# TCF-1 regulates the stem-like memory potential of HIV-specific CD8+ T cells in elite controllers

**DOI:** 10.1101/2020.01.07.894535

**Authors:** Rachel L. Rutishauser, Christian Deo T. Deguit, Joseph Hiatt, Franziska Blaeschke, Theodore L. Roth, Lynn Wang, Kyle Raymond, Carly E. Starke, Joseph C. Mudd, Wenxuan Chen, Carolyn Smullin, Rodrigo Matus-Nicodemos, Rebecca Hoh, Melissa Krone, Frederick M. Hecht, Christopher D. Pilcher, Jeffrey N. Martin, Richard A. Koup, Daniel C. Douek, Jason M. Brenchley, Rafick-Pierre Sékaly, Satish K. Pillai, Alexander Marson, Steven G. Deeks, Joseph M. McCune, Peter W. Hunt

**Author notes:** To whom correspondence should be addressed: Mailing Address: Zuckerberg San Francisco General Hospital 1001 Potrero Avenue, Building 3, Room 605 San Francisco, CA 94110.

## Abstract

Although many HIV cure strategies seek to expand HIV-specific CD8+ T cells to control the virus, all are likely to fail if cellular exhaustion is not prevented. A loss in stem-like memory properties (i.e., the ability to proliferate and generate secondary effector cells) is a key feature of exhaustion; little is known, however, about how these properties are regulated in human virus-specific CD8+ T cells. We found that virus-specific CD8+ T cells from humans and non-human primates naturally controlling HIV/SIV infection express more of the transcription factor, TCF-1, than non-controllers. HIV-specific CD8+ T cell TCF-1 expression correlated with memory marker expression and proliferative capacity and declined with antigenic stimulation. CRISPR-Cas9 editing of TCF-1 in human primary T cells demonstrated a direct role in regulating expansion capacity. Collectively, these data suggest that TCF-1 controls the stem-like memory properties of HIV-specific CD8+ T cells and provides a rationale for enhancing this pathway in T cell-based therapeutic strategies for HIV.

**One Sentence Summary:** TCF-1 is highly expressed in HIV-specific CD8+ T cells from elite controllers and directly regulates human CD8+ T cell expansion capacity in response to T cell receptor stimulation.

## Introduction

In the vast majority of individuals, infection with human immunodeficiency virus-1 (HIV) leads to chronic viremia that is not controlled by natural host immune responses. The failure of the immune system to control HIV is multifactorial, but one important component is the exhaustion of HIV-specific CD8+ T cells (*1–3*). Antigen-specific CD8+ T cell exhaustion occurs in the setting of chronic antigen exposure, and exhausted CD8+ T cells are defined by their impaired proliferation, effector cytokine production, and killing capacity after T cell receptor (TCR) stimulation with peptide (*4*). Therefore, even though they can initially control the viral load (*5–9*), HIV-specific CD8+ T cells that persist during the chronic phase of infection fail to eradicate infected cells from the body. Generating long-lived, non-exhausted antigen-specific CD8+ T cells that have stem-like T cell memory properties (i.e., the capacity to proliferate and to generate a burst of cytotoxic effector cells upon encountering cognate antigen) is the focus of several immunotherapeutic strategies (e.g., therapeutic vaccination, blockade of co-inhibitory receptors such as PD-1, and adoptive T cell therapies) for HIV (*10–12*) and other diseases in which CD8+ T cell exhaustion is observed, such as other chronic infections and cancers (*13*).

Unlike most HIV-infected individuals who have high viral loads in the absence of treatment with antiretroviral therapy (ART), roughly 1% of HIV-infected individuals, termed “controllers” here (also referred to as “elite controllers”), naturally control infection to undetectable levels in the blood in the absence of ART despite continuing to harbor HIV virus in lymphoid tissues throughout the body (*4*). Several lines of evidence (e.g., genomic associations between HIV control and peptide-binding domain sequence variants in MHC Class I alleles (*14–17*) and data from non-human primates infected with simian immunodeficiency virus [SIV] infection showing that controller animals experience viral rebound after CD8+ T cell depletion (*18*)) suggest that an effective HIV-specific CD8+ T cell response may contribute to the natural control of HIV in these individuals. Furthermore, compared to HIV-specific CD8+ T cells from either viremic or ART-suppressed non-controllers, HIV-specific CD8+ T cells isolated from controllers demonstrate stem-like memory T cell properties upon *in vitro* stimulation with peptide (*19–24*). Thus, controllers offer a model for understanding the mechanisms that support the generation of “functional” stem-like memory CD8+ T cell responses in humans in the context of a chronic infection. While several cell-intrinsic factors including the expression of PD-1 (*1, 2*) and other co-inhibitory receptors (*25–27*), activation of caspase-8 (*28*), and the expression of the transcription factor BATF (*29*) are known to regulate HIV-specific CD8+ T cell dysfunction, little is known about the molecular pathways that support the stem-like memory properties observed in HIV-specific CD8+ T cells from controllers.

In recent years, the Wnt-signaling transcription factor, TCF-1 (T cell factor 1; also known by its gene name, *TCF7*) (*30*), has been recognized as an important regulator of antigen-specific CD8+ T cell memory stem-like properties of proliferative and regenerative capacity (*31–33*). TCF-1 is highly expressed in naïve, central memory, and stem cell memory CD8+ T cells (*34*). In murine acute infection models, TCF-1 expression is required for the formation of long-lived antigen-specific memory responses that have the capacity to re-expand upon secondary challenge (*31–33, 35*). In the setting of chronic lymphocytic choriomeningitis (LCMV) infection in mice, TCF-1 marks the subpopulation of stem-like virus-specific CD8+ T cells that is capable of regenerating both the ongoing effector cell response as well as the proliferative burst of effector cells that is observed after blockade of the PD-1 signaling pathway (*36–38*). In humans, expression of TCF-1 in subpopulations of virus-specific CD8+ T cells has been reported in individuals chronically infected with hepatitis B virus (HBV) (*39*), hepatitis D virus (HDV) (*40*), or Epstein-Barr virus (EBV) (*40*), and TCF-1 expression has been correlated with virus-specific T cell expansion capacity in the context of infection with hepatitis C virus (HCV) (*37, 41*). Furthermore, in several recent clinical trials of individuals with melanoma, the size of the TCF-1-expressing subpopulation of intra-tumoral CD8+ T cells was strongly correlated with the clinical response to checkpoint blockade therapy (*42, 43*). At a mechanistic level, transcriptional programs supported by TCF-1 may counteract terminal differentiation and exhaustion programs mediated by other transcription factors (*36, 44*). In the context of HIV infection, TCF-1 expression has only been described in bulk CD8+ T cells that reside in the follicular regions of lymph nodes (*45*). To date, the expression of TCF-1 in HIV-specific CD8+ T cells and its relationship to functional capacity in these cells has not been described, nor has a causal relationship been established between TCF-1 and stem-like memory properties in human virus-specific T cells. Here, we uncover key molecular differences in HIV-specific CD8+ T cells from HIV controllers and non-controllers by exploring the relationship between TCF-1 expression and HIV control and demonstrating a causal role for this transcription factor in retaining stem-like memory properties in HIV-specific CD8+ T cells in humans.

## Results

### HIV- and SIV-specific CD8+ T cells from natural controllers have high TCF-1 expression

We first sought to confirm in our cohort the observation that HIV-specific CD8+ T cells from individuals who naturally control HIV infection to undetectable levels in the blood off of ART (“controllers”) have enhanced proliferative capacity (*19, 21, 28*). To do this, we identified HLA-typed participants with HIV enrolled in the San Francisco-based SCOPE cohort who had detectable HIV-specific CD8+ T cell responses in the peripheral blood by MHC Class I multimer staining (“multimer+”). Participants were sampled from one of three clinical groups: controller, viremic, or ART-suppressed (n=12, 13 and 10 participants in each group, respectively; see Materials and Methods for definitions, Fig. S1 for multimer gating, Fig. 1A for multimer+ cell frequencies in the different groups). We labeled peripheral blood mononuclear cells (PBMCs) from these individuals with the cell proliferation dye, CellTrace Violet (CTV), and stimulated them *in vitro* for six days with peptide pools containing the peptide recognized by the multimer+ CD8+ T cell population. As others have reported, we found that the multimer+ HIV-specific CD8+ T cells from controllers proliferated more robustly and tended to demonstrate a greater absolute increase in the proportion of Granzyme B-expressing cytotoxic effector cells after stimulation compared to multimer+ cells from viremic or ART-suppressed individuals (Fig. 1B, Fig. S2). We also noted that the enhanced proliferative capacity of HIV-specific CD8+ T cells from controllers did not appear to be explained by an increased frequency of cells with a central memory (TCM) phenotype (CD45RA-CCR7+; Fig. 1C) (*46*). Rather, multimer+ cells evaluated directly *ex vivo* across clinical groups tended to fall into a transitional memory phenotype (TTM: CD45RA+CCR7-CD27+) regardless of the presence or mechanism of viral control.

**Fig. 1.**
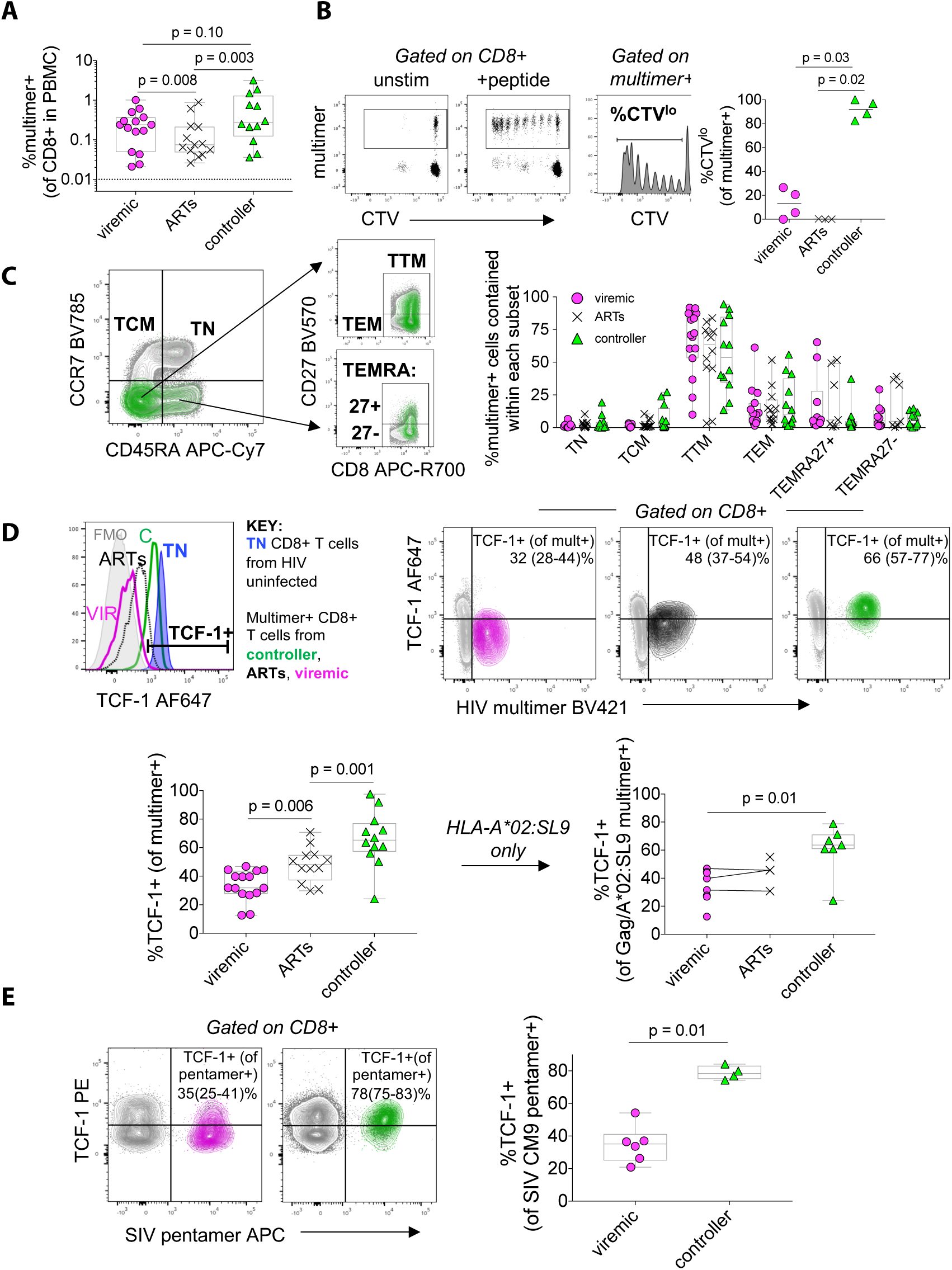
TCF-1 expression is elevated in HIV- and SIV-specific CD8+ T cells from controllers. (**A**) Frequency of peripheral blood multimer+ HIV-specific CD8+ T cells. (**B**) Proliferation of HIV-specific CD8+ T cells in response to six-day *in vitro* cognate peptide stimulation as measured by dilution of cell-trace violet (CTV). (**C**) Gating strategy (left; green: multimer+ from controller; grey: bulk CD8+ T cells) and distribution (right) of effector-memory phenotypes amongst multimer+ cells (TN, naïve; TCM, central memory; TTM, transitional memory; TEM, effector memory; TEMRA, effector memory-RA). (**D**) Gating (top left; TCF-1+ population gated based on CD8+ TN population from an HIV-uninfected participant, blue), representative flow plots (top right; median (range)), and summary data (bottom) showing TCF-1 expression in multimer+ HIV-specific CD8+ T cells from viremic (“VIR”; magenta), ART-suppressed (“ARTs”; black), and controller (“C”; green) individuals of all multimer specificities (left) and within the HIV Gag/HLA-A*02:SL9 multimer+ population (right). (**E**) TCF-1 expression in the SIV Gag/Mamu-A*01:CM9 multimer+ population from viremic and controller macaques. Phenotypes assessed by flow cytometry. FMO, fluorescence-minus-one control. Mult, multimer. Box plots: median ± interquartile range. Statistical testing: Linear mixed effects models to account for clustering within participants (A, C, D), Kruskal-Wallis followed by Dunn’s multiple comparison testing (B), Wilcoxon Rank Sum (E).

Given the association between high TCF-1 expression and the regenerative capacity of antigen-specific CD8+ T cells in mice (*32, 35*), we hypothesized that this transcription factor would be more highly expressed in HIV-specific CD8+ T cells from controllers. Indeed, we found that the proportion of multimer+ HIV-specific CD8+ T cells expressing TCF-1 was highest in controllers, followed by ART-suppressed, and then viremic participants (median 62% vs. 51% vs. 35% TCF-1+; p=0.006 viremic vs. ART-suppressed, p=0.001 ART-suppressed vs. controller; Fig. 1D). This observation held true even when we restricted our analysis to a single multimer response (HIV-specific CD8+ T cells that recognize the Gag-derived peptide SLYNTVATL presented in the context of HLA-A*02). Compared to viremic individuals, controllers also had a higher frequency of TCF-1+ cells within the predominant TTM and TEM subsets of multimer+ cells (Fig. S3), confirming that the higher expression of TCF-1 in multimer+ cells from controllers is not simply explained by differential memory subset distribution.

Finally, rhesus macaques (RM) that control simian immunodeficiency virus (SIV) in the absence of therapy also have highly functional, relatively non-exhausted SIV-specific CD8+ T cells (*47*). We found that ART-naïve controller macaques with low viral loads (< 1,000 copies/mL) had higher expression of TCF-1 in their Mamu-A*01-restricted CM9 SIV-specific CD8+ T cells compared to untreated animals with high viral loads (median 290,500 copies/mL; p=0.01; Fig. 1E and Table S2). Therefore, across two primate species, natural control of chronic retroviral infection is associated with the generation of a virus-specific CD8+ T cell population with high TCF-1 expression.

### TCF-1-expressing HIV-specific CD8+ T cells are phenotypically less differentiated

Although we found that the distribution of classic effector-memory phenotypes within the HIV-specific CD8+ T cell population is not affected by presence or mechanism of viral control (Fig. 1C), multimer+ CD8+ T cells evaluated directly *ex vivo* from HIV controllers had several other phenotypic features more consistent with resting central memory cells: a larger proportion expressed CD127 while a smaller proportion expressed the effector protein Granzyme B or high levels of the effector differentiation transcription factor, T-bet (Fig. 2A). In contrast, viremic participants tended to have a more effector/effector-memory-like phenotype of multimer+ CD8+ T cells. Eomesodermin, another T-box transcription factor important for memory differentiation (*48, 49*), did not appear to be differently expressed in HIV-specific CD8+ T cells from controllers (Fig. S4A). Similar to humans, SIV-specific CD8+ T cells from controller compared to typical progressor macaques with high viral loads had a more resting memory-like phenotype (i.e., they had a higher expression of CD127 and LEF-1, a transcription factor that can coordinate with TCF-1 to activate Wnt signaling (*30, 35, 50*), and a trend towards lower expression of Granzyme B) and they appeared to be less activated (with lower CD69 expression and a trend towards a smaller proportion of Ki-67+ cells; Fig. S4B). Furthermore, the expression of the chemokine receptor, CXCR5, was higher in TCF-1+ compared to TCF-1-SIV-specific CD8+ T cells amongst viremic animals (p=0.03). This relationship was also observed in the four controller animals with a trend (p=0.125) towards significance (Fig. S4B).

**Figure 2.**
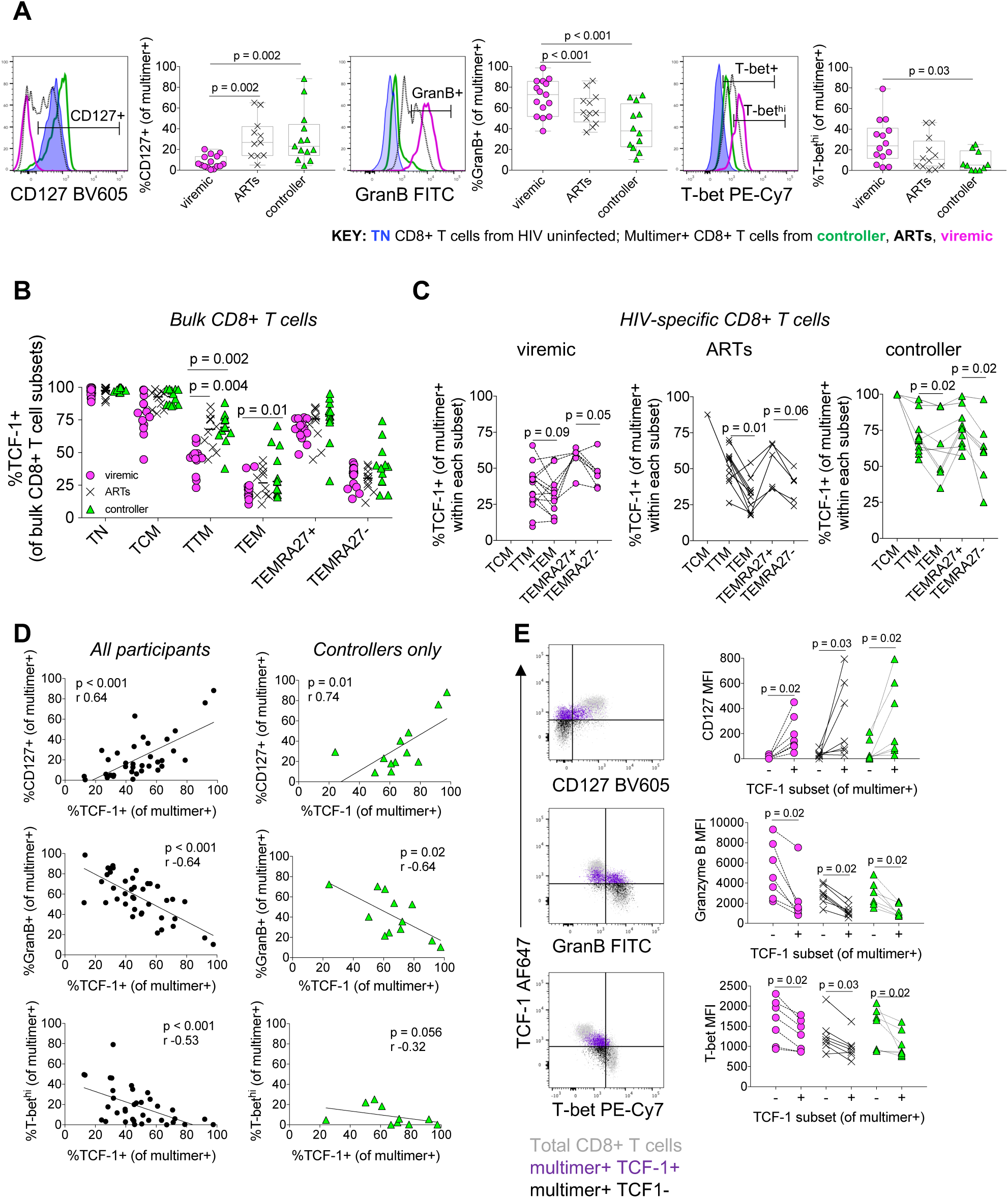
High TCF-1 expression in HIV-specific CD8+ T cells is associated with a memory-like phenotype. (**A**) Phenotype of HIV-specific multimer+ CD8+ T cells from viremic, ART-suppressed, or controller individuals (compared to naïve CD8+ T cells from an HIV negative individual). TCF-1 expression is higher in less differentiated naïve and effector-memory subsets of bulk (**B**) and HIV-specific (**C**) CD8+ T cells. (**D**) Correlation between the expression of TCF-1 and other phenotypic markers in multimer+ CD8+ T cells (GranB, Granzyme B). (**E**) Expression of CD127, Granzyme B, and T-bet within TCF-1+ versus TCF-1-HIV-specific CD8+ T cells. Statistical testing: Linear mixed effects models to account for clustering within participants (A, B), Wilcoxon Signed Rank test (C, E), Spearman correlation (D).

In adults without HIV infection, TCF-1 expression has been found to be higher in less-differentiated bulk CD8+ T cell subsets (i.e., TN and TCM; (*51*)). We observed a similar pattern in our cohort of participants with HIV in both bulk (Fig. 2B) and HIV-specific CD8+ T cells (Fig. 2C). In particular, CD8+ T cell subsets that express CD27, a tumor necrosis factor family protein important for T cell survival (*52, 53*) (i.e., TN, TCM, TTM, and CD27+ TEMRA cells) tended to express more TCF-1 compared to subsets that do not express CD27 (i.e., TEM and CD27-TEMRA cells). Furthermore, mirroring the antigen-specific population, controllers also had higher TCF-1 expression within the gated TTM and TEM bulk CD8+ T cell subsets compared to viremic individuals (Fig. 2B).

Within the population of multimer+ HIV-specific CD8+ T cells, we found that TCF-1 expression was associated with patterns of phenotypic marker expression typically found in central memory CD8+ T cells, with a positive correlation between TCF-1 and CD127 expression (Spearman correlation, r=0.64, p<0.0001), and a negative correlation between TCF-1 and Granzyme B (r=-0.64, p<0.001), and T-bet expression (r=-0.53, p<0.001), both across the whole cohort and specifically within HIV-specific CD8+ T cells isolated from controllers (Fig. 2D). Finally, TCF-1-expressing HIV-specific CD8+ T cells in all three groups expressed significantly more CD127, and less Granzyme B and T-bet (Fig. 2E), as well as more of the memory-associated transcription factor FOXO1 (per-cell total protein level; Fig. S4C).

### Varied co-inhibitory receptor expression on HIV-specific CD8+ T cells from controllers

A number of cell surface co-inhibitory receptors (e.g., PD-1, TIGIT, CD160, and 2B4) are upregulated in the setting of chronic T cell receptor stimulation and their expression has been associated with functional exhaustion of antigen-specific CD8+ T cells in HIV disease (*1, 2, 25, 54–56*) and other settings (*13*). Consistent with their observed robust capacity to proliferate and generate secondary effector cells in response to stimulation, HIV-specific CD8+ T cells from controllers expressed PD-1 less frequently than those from both viremic and ART-suppressed individuals (Fig. 3A). Although some controllers had a large subpopulation of multimer+ CD8+ T cells that expressed some PD-1, the absolute level of expression of PD-1 on multimer+ cells from controllers was low (as measured by the median fluorescence intensity, MFI, as well as the proportion of HIV-specific CD8+ T cells that express high levels of PD-1, “PD-1^hi^”; Fig. 3A). Across the cohort, the proportion of HIV-specific CD8+ T cells that expressed TCF-1 negatively correlated with the proportion expressing PD-1 and the directionality (but not significance) of this relationship was also maintained when evaluating controllers alone (Fig. 3B). TCF-1-expressing cells from viremic individuals have lower expression of PD-1 compared to TCF-1-negative cells; this relationship, however, did not appear to hold amongst HIV-specific CD8+ T cells from ART-suppressed or controller individuals (Fig. 3C). Interestingly, unlike true memory cells generated in the context of acute infection, HIV-specific CD8+ T cells from controllers compared to ART-suppressed or viremic individuals did not express lower levels of three other co-receptors that have been associated with antigen-specific CD8+ T cell functional exhaustion (TIGIT, CD160, 2B4; Fig. 3D).

**Figure 3.**
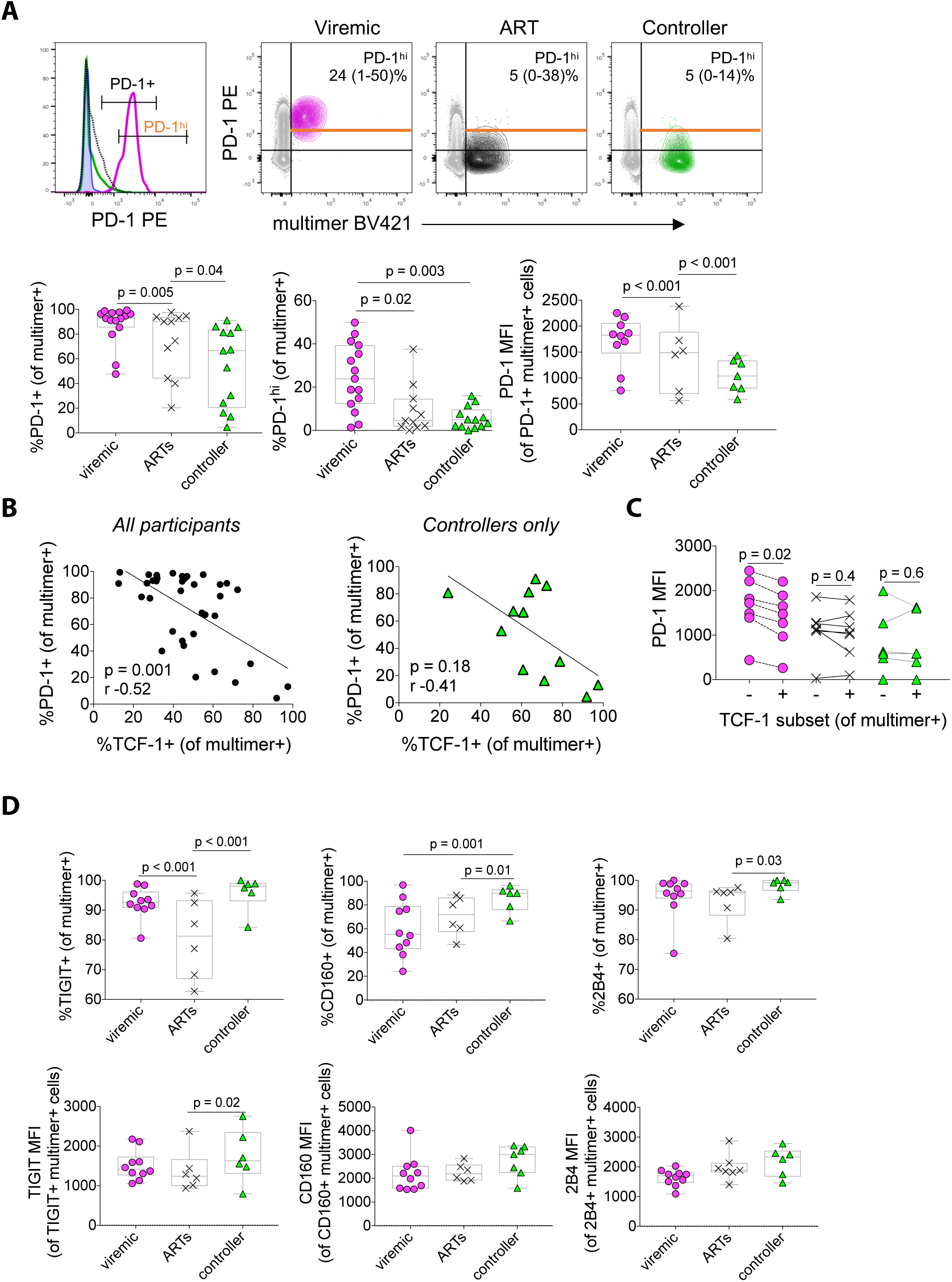
HIV-specific CD8+ T cells have lower expression of PD-1 but not other co-inhibitory receptors. (**A**) Expression of PD-1 in multimer+ HIV-specific multimer+ CD8+ T cells (annotated with median and range; MFI, median fluorescence intensity). (**B**) Correlation between PD-1 and TCF-1 expression in multimer+ HIV-specific CD8+ T cells. (**C**) Expression of PD-1 within TCF-1+ versus TCF-1-HIV-specific CD8+ T cells. (**D**) Expression of TIGIT, CD160, and 2B4 in multimer+ HIV-specific multimer+ CD8+ T cells. Statistical testing: Linear mixed effects models to account for clustering within participants (A, D), Wilcoxon Signed Rank (C), Spearman correlation (B).

### TCF-1 expression in HIV-specific CD8+ T cells is downregulated by T cell receptor stimulation

Because low TCF-1 expression is associated with CD8+ T cells that are more differentiated towards an effector fate, we asked whether TCF-1 expression is reduced upon stimulation of HIV-specific CD8+ T cells with their cognate peptide to induce secondary effector differentiation. After six-day *in vitro* peptide stimulation, the proportion of HIV-specific CD8+ T cells expressing TCF-1 decreased in the undivided (CTV^hi^) versus the divided (CTV^lo^) population from a median 44% to 1% (p < 0.001; Fig. 4A). In a sample where clear division peaks were discernable, this reduction appeared to mostly take place over the course of the first three divisions and occurred inversely to the acquisition of secondary effector characteristics (e.g., the expression of Granzyme B and Perforin; Fig. 4B).

**Figure 4.**
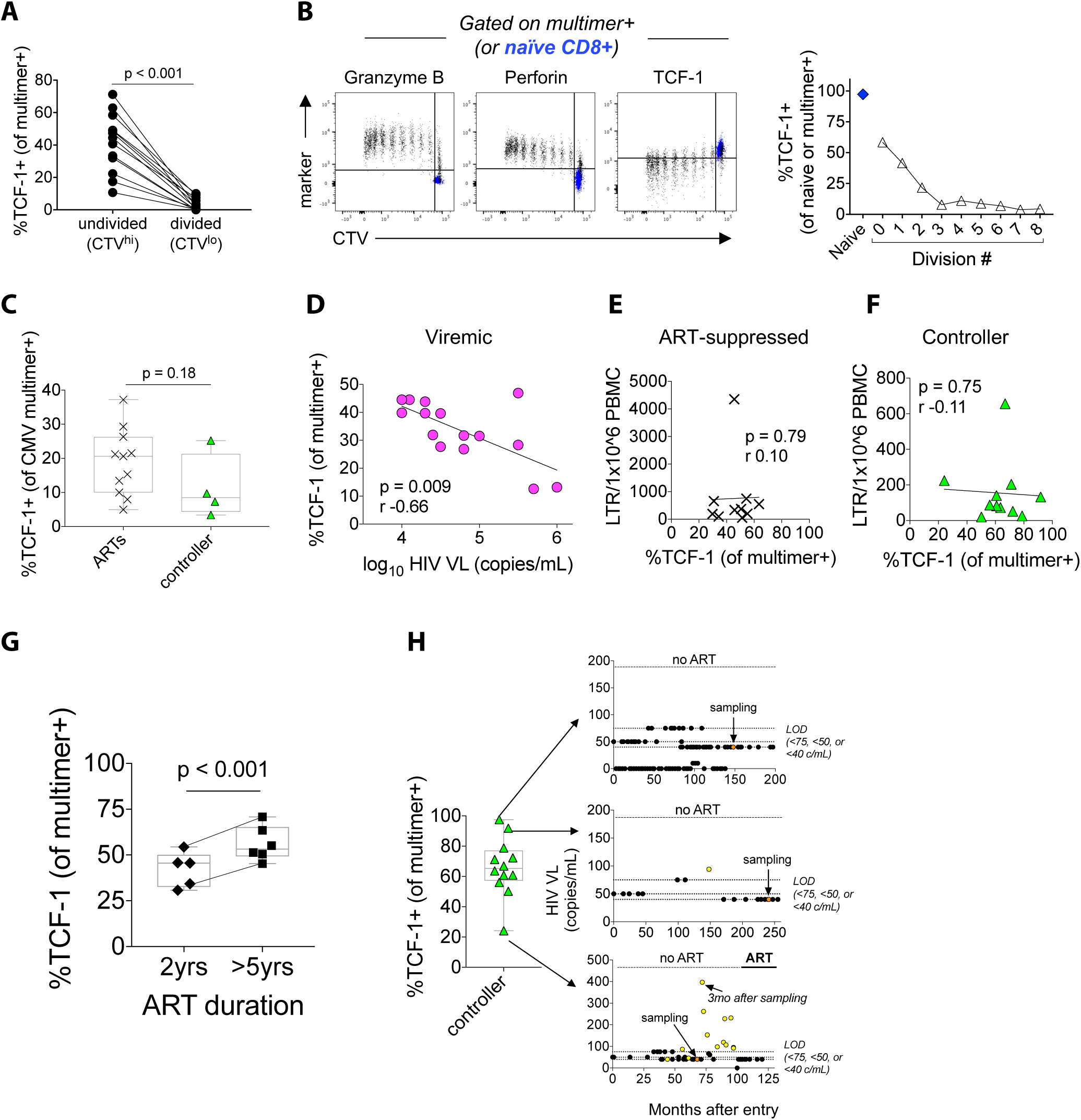
TCF-1 expression is negatively correlated with antigen exposure. (**A**) TCF-1 expression in HIV-specific CD8+ T cells after 6 days of *in vitro* peptide stimulation in divided versus undivided cells. (**B**) TCF-1 expression in multimer+ CD8+ T cells (black) in each division peak after 6 days of *in vitro* peptide stimulation (naïve CD8+ T cells shown in blue for reference). (**C**) TCF-1 expression in multimer+ CMV-specific CD8+ T cells from HIV-infected individuals (ART-suppressed versus controller). (**D**) Negative correlation between TCF-1 expression and plasma HIV viral load (VL). Relationship between TCF-1 expression in HIV-specific CD8+ T cells and HIV DNA levels (long-term repeat [LTR] copies per million PBMC) in ART-suppressed (**E**) and controller (**F**) individuals. (**G**) Expression of TCF-1 in HIV-specific CD8+ T cells in ART-suppressed individuals, depending on the duration of ART. (**H**) Timing of PBMC in HIV elite controllers with the highest (right panels, top two participants) and lowest (bottom participant) levels of TCF-1 in HIV-specific CD8+ T cells. Detectable HIV viral load measurements noted in yellow. LOD=limit of detection. c/mL=copies/mL. Statistical testing: Wilcoxon Rank Sum (C), Spearman correlation (D-F), Linear mixed effects models to account for clustering within participants (G).

We next explored how TCF-1 expression in antigen-specific CD8+ T cells relates to the level of stimulation they receive from ongoing chronic infection *in vivo*. We first asked whether the level of TCF-1 in antigen-specific CD8+ T cells that recognize a different virus (cytomegalovirus, CMV) is affected by the non-antigen-specific physiologic differences present between the HIV clinical groups (e.g., differences in the levels of cytokines and other inflammatory mediators) (*57*). We observed a similar level of TCF-1 expression between CMV-specific CD8+ T cells observed from ART-suppressed versus controller individuals (Fig. 4C). We then asked how different levels of antigen stimulation *in vivo* relate to TCF-1 expression in HIV-specific CD8+ T cells. Amongst viremic participants, higher TCF-1 expression multimer+ HIV-specific CD8+ T cells correlated with lower HIV viral loads (r=-0.66, p=0.009; Fig. 4D). The directionality (but not the significance) of this relationship remained in the subset of participants (n=9) with Gag/A*02:SL9 responses (r=-0.40, p=0.28; Fig. S5), despite the fact that the dominant plasma viral species from all seven of the participants for whom we have plasma viral sequence data available demonstrated evidence of viral evolution and CD8+ T cell epitope escape at this peptide sequence at the time of sampling (Table S4).

In contrast to the negative correlation between HIV-specific CD8+ T cell TCF-1 expression and viral load in viremic participants, we did not observe a significant relationship between HIV cell-associated DNA levels in PBMCs and TCF-1 expression in either ART-suppressed individuals or elite controllers (Fig. 4E and F). Interestingly, a longer duration of ART (>5 years) was associated with a higher level of TCF-1 expression in HIV-specific CD8+ T cells (Fig. 4G; p<0.001). Of the controllers we included in our study, we noted that one individual had a particularly low frequency of TCF-1-expressing multimer+ cells (24%) compared to the rest of the controller group (median 67%; Fig. 1D, also shown in Fig. 4H). Interestingly, this individual had multiple detectable viral load measurements of 100-300 copies/mL starting three months after the PBMC sample date (Fig. 4G, right side, bottom). In contrast, the two controller individuals with the largest proportion of TCF-1-expressing multimer+ cells in the cohort (Fig. 4G, right side, top and middle) have continued to maintain HIV suppression with undetectable viral loads in the absence of ART and with frequent testing for 16 and 20 years in the cohort, with the exception of a single detectable viral load (< 100 copies/mL) more than eight years prior to sampling in one individual.

### TCF-1 regulates human CD8+ T cell expansion capacity

We found that TCF-1 expression is associated with a T cell memory-like phenotype in HIV-specific CD8+ T cells. We next asked whether TCF-1 expression is correlated with the stem-like memory property of expansion capacity in HIV-specific CD8+ T cells. Indeed, we found that the level of TCF-1 expression in multimer+ HIV-specific CD8+ T cells directly *ex vivo* correlated with their ability to proliferate in response to six-day *in vitro* stimulation with cognate peptide (r=0.88, p=0.007; Fig. 5A). We also found that multimer+ CMV-specific CD8+ T cells with higher TCF-1 expression demonstrate a trend towards increased expansion after cognate peptide stimulation (r=0.53, p=0.054; Fig. 5B).

**Figure 5.**
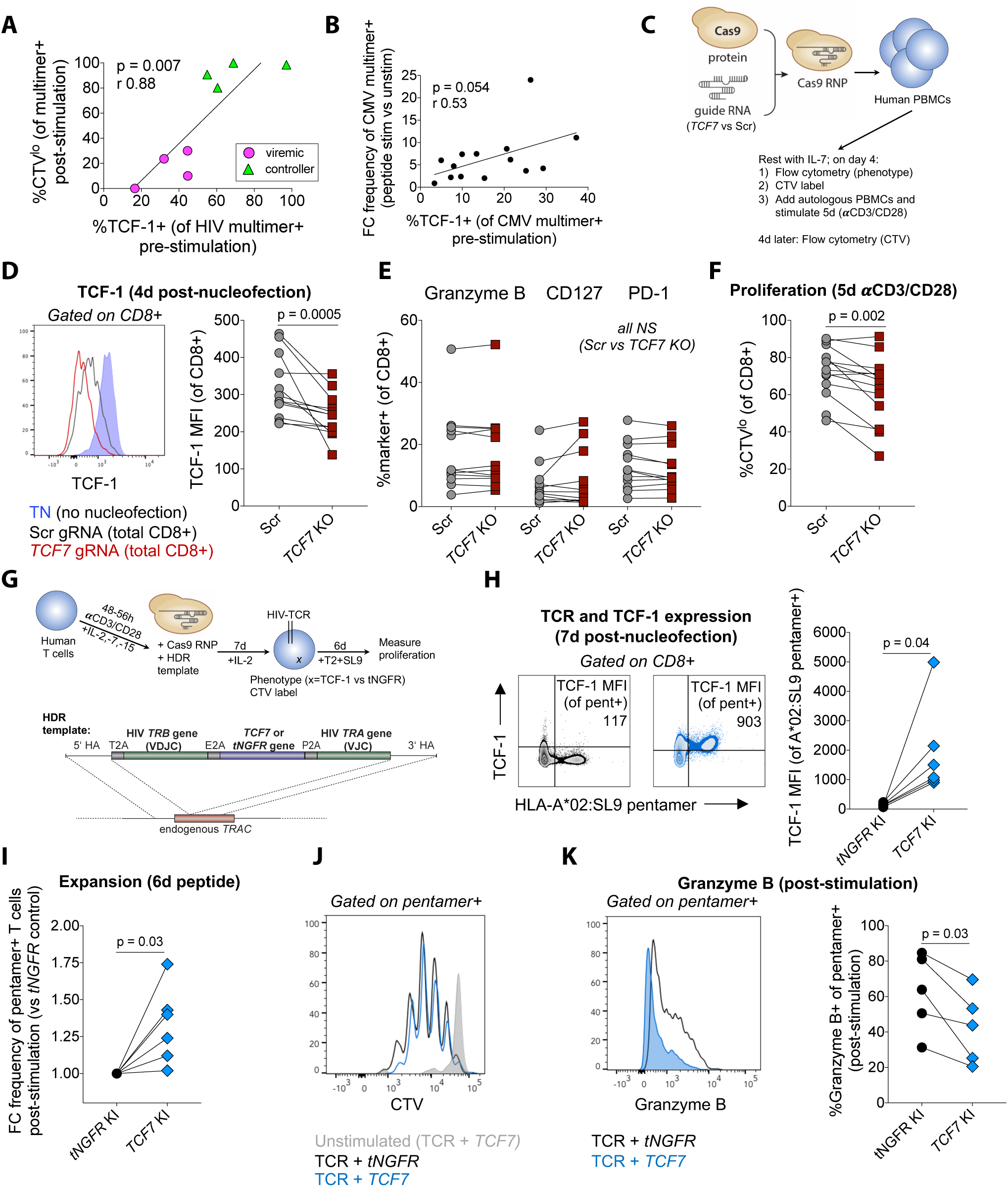
TCF-1 directly regulates human CD8+ T cell responses to TCR stimulation. (**A**) Correlation between TCF-1 expression in HIV-specific CD8+ T cells and the proportion that divide after six-day *in vitro* peptide stimulation. (**B**) Correlation between TCF-1 expression in CMV-specific CD8+ T cells and their expansion after six-day *in vitro* peptide stimulation (fold change [FC] in the frequency of CMV-specific CD8+ T cells [gated on total CD8+ T cells], stimulated versus unstimulated cells). (**C**) CRISPR-Cas9-mediated deletion of *TCF7* (Scr, scrambled guide (g)RNA; RNP, ribonucleoprotein). (**D**) TCF-1 protein downregulation in CD8+ T cells after *TCF7* knockout (KO; red) compared to electroporation with Scr gRNA (grey). (**E**) CD8+ T cell phenotype after *TCF7* KO. (**F**) Proportion of divided CD8+ T cells after five-day stimulation with αCD3/CD28. (**G**) Generation of HIV-specific T cell receptor (TCR)-T cells using CRISPR-Cas9 knock-in (KI) of HIV-specific TCR and *TCF7* (versus truncated Nerve Growth Factor Receptor [*tNGFR*]). (**H**) TCF-1 protein expression after *tNGFR* (black) or *TCF7* (blue) KI. (**I**) Frequency of *TCF7* KI TCR-T cells after six-day *in vitro* stimulation with SL9 peptide loaded on the T2 cell line (FC versus *tNGFR* KI). (**J**) CTV tracings of TCR-T cells (unstimulated or stimulated). (**K**) Granzyme B expression in *TCF7* KI TCR-T cells. Statistical testing: Spearman correlation (A, B), Wilcoxon Signed Rank (D, E, F, H, I, K).

In order to assess whether TCF-1 directly regulates human CD8+ T cell proliferative capacity, we employed CRISPR-Cas9-targeted genomic editing approaches to knock out or overexpress the *TCF7* gene in primary human T cells. We first asked whether genetic ablation of *TCF7* impairs proliferation. In comparison to cells electroporated with Cas9 ribonucleoproteins (RNPs) with a control (scrambled) guide RNA not predicted to cut any site in the human genome, CD8+ T cells in which *TCF7* was targeted by Cas9 RNPs showed no change in several phenotypic markers but had impaired proliferative responses after five-day polyclonal T cell receptor (TCR) stimulation with αCD3/CD28 antibodies (p=0.002; Fig. 5C-F). Using a recently-developed non-viral gene knock-in approach, we genetically modified primary human T cells to replace the endogenous TCR with an HIV-specific TCR co-expressed with TCF-1 or an irrelevant control protein, truncated Nerve Growth Factor Receptor (tNGFR), both under the control of the endogenous TCRα gene (*TRAC*) promoter (*58, 59*) (Fig. 5G-H). Six-day *in vitro* stimulation of these cells with cognate peptide-loaded antigen presenting cells led to an increase in the recovery of expanded TCR-T cells that overexpress the TCF-1 protein compared to TCR-T cells expressing tNGFR (p=0.03; Fig. 5I). Based on CTV tracings, the increased recovery of the *TCF7* knock-in TCR-T cells appeared to be due to enhanced survival of the proliferated cells rather than an absolute increase in the proportion of cells that proliferated (Figs. 5J and S6). Consistent with its role as a regulator of stem-like memory properties, over-expression of TCF-1 was also associated with a small but significant decrease in the expression of the cytolytic protein, Granzyme B, in HIV-specific TCR-T cells (Fig. 5K).

## Discussion

The goal of many immune-based strategies that aim to enhance the control of HIV is to elicit functional, non-exhausted, and durable HIV-specific CD8+ T cells that have the stem-like memory T cell capacity to rapidly expand into secondary effector cells that can kill HIV-infected cells (*10, 12*). We hypothesized that TCF-1, a Wnt signaling transcription factor known to regulate T cell “stemness” in several other contexts, might also regulate the expansion capacity of HIV-specific CD8+ T cells. We found that MHC Class I multimer+ HIV-specific CD8+ T cells with high proliferative capacity from HIV controllers compared to cells from non-controllers on or off of ART have a larger TCF-1+ subpopulation, that TCF-1 expression in these cells is associated with higher expression of other proteins associated with T cell memory function and lower PD-1 expression, that knockout of the *TCF7* gene impairs polyclonal CD8+ T cell proliferation, and that *TCF7* gene overexpression in genetically engineered HIV-specific CD8+ T cells enhances their accumulation after peptide stimulation. Identifying the pathways that support antigen-specific CD8+ T cell stem-like capacity in the context of HIV is critical not only to directly inform the development of improved CD8+ T cell-based therapies to control HIV, but also to more broadly understand how this unique differentiation state is regulated in human CD8+ T cells.

These data are the first demonstration, to our knowledge, of a relationship between TCF-1 expression in HIV-specific CD8+ T cells and elite controller status, and also the first indication of a direct role for *TCF7*/TCF-1 in the regulation of expansion capacity in human virus-specific T cells. Our phenotypic findings align with several studies that have demonstrated a role for TCF-1 in supporting memory cell capacity and countering terminal effector differentiation and exhaustion programs in mice with acute and chronic viral infections, in human chronic hepatitis C infection, and in mouse and human cancers (*31, 32, 36–38, 41, 44, 60–64*). Taken together with our functional studies, these data provide a clear rationale for further exploration of this pathway as a potential target for modulation in T cell-based therapies for HIV. For example, therapeutic vaccine regimens (*10*) might be improved by employing adjuvants that promote TCF-1 expression in vaccine-elicited HIV-specific CD8+ T cells. Similarly, overexpressing TCF-1 in HIV-specific receptor-engineered T cells prepared for adoptive transfer (either TCR-T cells, as we have made here, or chimeric antigen receptor [CAR] T cells (*65*)) might be one approach to overcoming the major challenge of generating T cells that can persist long-term and avoid exhaustion after *in vivo* infusion (*66*).

Identifying a mechanistic pathway that supports the stem-like memory T cell functional capacity of HIV-specific CD8+ T cells in individuals who naturally control this chronic infection raises two important questions. First, is this capacity a cause or a consequence of viral control in these individuals? Because TCF-1 expression decreases in antigen-specific CD8+ T cells upon TCR stimulation, it is possible that the higher levels of TCF-1 that we observe in controllers is at least in part a consequence of low levels of antigen exposure in these individuals. Interestingly, we observed low TCF-1 expression in HIV-specific CD8+ T cells from viremic non-controllers even after either CD8+ T cell epitope escape or viral suppression with antiretroviral therapy. These results suggest that there may be a threshold of high antigen exposure beyond which, even if there is subsequently a reduction in TCR stimulation, antigen-specific CD8+ T cells are unable to recover stem-like memory properties. At the same time, while it is difficult to mechanistically show this in humans, it is also very plausible that the enhanced ability of HIV-specific CD8+ T cells from controllers to mount a robust proliferative response (a capacity that is directly linked to their *in vitro* cytotoxicity) is causally linked to their ability to naturally contain the virus.

A second question raised by our data is whether promoting stem-like memory T cell capacity is the optimal differentiation state to target for T cell-based interventions to treat HIV, a setting in which the therapy-derived T cells must outpace viral rebound after antiretroviral therapy discontinuation presumably by (1) being present in high enough numbers, (2) being localized in the correct tissues, and (3) having sufficient (and appropriate) effector functions to eliminate infected cells. TCF-1 is a promising target to improve T cell-based HIV therapies because animal models suggest that TCF-1 regulated stem-like properties are essential for protective CD8+ T cell secondary recall responses after natural infection or vaccination (*31, 35*) and the survival of CD8+ T cells that experience chronic stimulation (*37, 38*). However, our data in TCR-T cells genetically engineered to overexpress *TCF7* also support findings in other studies showing that constitutive TCF-1 expression or activation of the Wnt signaling pathway promotes T cell “stemness” at the expense of effector differentiation (*33, 67*). In other words, permanent over-expression of TCF-1 might support the proliferative capacity of HIV-specific CD8+ T cells such that they would be able to expand robustly and survive better upon secondary antigen encounter; however, without the ability to downregulate TCF-1, the daughter cells would potentially lack sufficient cytotoxic effector functions. Even if a therapeutic intervention were designed to generate TCF-1+ T cells that had the ability to downregulate TCF-1 and differentiate into effector cells upon antigenic stimulation, because TCF-1+ T cells themselves are not effector-differentiated or poised for immediate killing, it is possible that their delayed responsiveness would be outpaced by viral rebound.

In addition to these questions, for the development of therapeutic vaccination and adoptive T cell therapies for HIV, it will be essential to understand how targeting the TCF-1 pathway in human T cells affects the formation of tissue resident memory (TRM) and follicular cytotoxic (Tfc) CD8+ T cells. It has been proposed that CD8+ T cells that occupy these differentiation states would provide a rapid response to nearby reactivated virus because they are localized closer to the sites of HIV reservoir persistence within lymphoid tissues (and, in particular, B cell follicles; (*68–70*)). Indeed, studies have suggested that lymphoid tissue from humans and non-human primates that naturally control retroviral infection is enriched for CD8+ T cells that express the TRM-associated protein, CD69 (*71, 86*), as well as CD8+ T cells that express the Tfc-associated chemokine receptor, CXCR5 (*72, 73*). Studies in mice suggest that TCF-1 may directly promote the generation of Tfc (*38*) but inhibit the formation of TRMs (*74*). Although little is known about the relationship between TCF-1 and the formation of these subsets in humans, in our study we did observe a correlation between the expression of TCF-1 and CXCR5 in SIV-specific CD8+ T cells isolated from the spleens of infected rhesus macaques. To effectively harness this pathway for T cell therapies for HIV and other diseases such as cancer, it will be important to understand the direct impact of TCF-1 expression on TRM/Tfc differentiation as well as other T cell migratory properties.

As a mostly cross-sectional study performed on peripheral blood samples in humans and splenocytes from non-human primates, our study has some limitations. First, because we have not sampled other tissues, we cannot extrapolate our conclusions to infer how TCF-1 interacts with other cellular phenotypes or functions in tissues. Based on work by other groups, we anticipate that HIV-specific CD8+ T cells isolated from lymph nodes (*71, 75*) and the gut (*76, 77*) will have substantially different phenotypes compared to those in peripheral blood and it would be important to assess TCF-1 expression in these tissues in follow-up studies. Second, due to inherent limitations in the CRISPR editing protocols, our functional experiments did not allow us to directly test the role of TCF-1 in regulating stem-like memory capacity in endogenous HIV-specific CD8+ T cells. However, as protocols in this field are rapidly developing, we hope we will be able to address this question in the future.

Our study provides important rationale to support further investigations into the fundamental biology of TCF-1 activity in human T cells and how to harness this pathway to optimize T cell-based therapies for HIV. Although downstream transcriptional and epigenetic targets of TCF-1 have been studied in developing thymocytes and other immune cell types in mice (*44*), the pathways via which TCF-1 exerts its effects on virus-specific T cells (in mice or humans) are currently unknown. In therapeutic settings, approaches to overexpress TCF-1 in a manner such that levels can be modulated to allow for T cell differentiation while preserving the positive influence of TCF-1 on T cell longevity should be explored. Finally, it will be critical to study whether high levels of TCF-1 in vaccine-elicited T cells correlate with the long-lasting durability of the response, affect the localization of the antigen-specific T cells, and, ultimately promote the ability of a therapeutic vaccine regimen to control viral rebound after antiretroviral treatment discontinuation.

## Materials and Methods

### Study Design

Our studies of human samples from individuals infected with HIV were designed as an analysis of retrospectively-collected peripheral blood mononuclear cells (PBMCs) from individuals enrolled in an existing cohort. We selected samples for inclusion in the study based on screening cohort participants for MHC Class I multimer+ HIV-specific CD8+ T cell responses and did not perform a power analysis prior to selecting samples for inclusion. Flow cytometry samples were excluded if viability was < 80%. No blinding was performed. Number of participants included and number of experiments performed for each assay is indicated below.

### Human Study Participants and Samples

This study sampled PBMCs retrospectively collected from HIV-infected participants enrolled in the Zuckerberg San Francisco General Hospital clinic-based SCOPE cohort (and individuals co-enrolled in the related Options cohort) who had a documented positive HIV antibody test, who were not on ART at the time of enrollment, and who had HIV-specific CD8+ T cell responses detectable by MHC Class I multimer staining (Table S1, and see below for multimer specificities and staining). Individuals were classified into one of three groups: (1) viremic (plasma viral load [VL] > 9,000 copies/mL; infected for at least six months [and greater than two years for all but one participant] prior to sampling; most individuals were ART-naïve, but if not, they were off ART for at least two years); (2) ART-suppressed (VL < 40 copies/mL; infected and untreated for at least two years, and then on ART with suppressed VL for at least two years prior to sampling); (3) controller (VL < 100 copies/mL; infected for at least two years, with no viral loads > 200 copies/mL for at least three months prior to and after sampling; most individuals were ART-naïve, but if not, they were off ART for at least two years). CD4+ T cell counts and HIV-1 plasma RNA levels were measured in all cohort participants by CLIA-certified clinical laboratory assays at the time of entry into the cohort and approximately every 3-4 months thereafter. This study also sampled de-identified PBMCs from blood donors known to have negative testing for HIV, HCV, and HBV (Stanford Blood Bank and Vitalant). Peripheral blood mononuclear cells (PBMCs) were collected and cryopreserved as described previously (*78, 79*). The UCSF Committee on Human Research approved this study, and participants gave informed, written consent before enrollment.

### Rhesus Macaque Characteristics, SIV Infection, and Rhesus Splenocyte Flow Cytometry

Splenic mononuclear cell suspensions were obtained from 10 *Mamu-A*01+* rhesus macaques (*Macaca mulatta*) infected with either SIVmac239, SIVsmE543, or SIVsmE660 and with established SIV infection or AIDS. Macaques were classified as viremic (> 1,000 copies/mL; n=6) or controller (< 1,000 copies/mL; n=4) based on plasma RNA viral loads (Table S2). All experimental procedures were approved by the National Institute of Allergy and Infectious Diseases Division of Intramural Research Animal Care and Use Program as part of the National Institutes of Health Intramural Research Program (protocols LMM6 and LVD26). Rhesus macaques were housed and sustained in accordance with standards established by the Association for Assessment and Accreditation of Laboratory Animal Care (AAALAC). Cryopreserved samples were homogenized as described previously (*80*). Tissue homogenates were washed twice in RPMI 1640 medium supplemented with 10% fetal bovine serum, 2 mM L-glutamine, and 1% penicillin/streptomycin (HyClone via GE Healthcare Life Sciences, Pittsburgh, PA, USA). SIV-specific CD8+ T cells were identified using Mamu-A*01-CTPYDINQM Gag181-189 (CM9) Pro5 MHC Class I Pentamers (ProImmune, Oxford, England). Cells were stained with the viability marker LIVE/DEAD Aqua (Thermo Fisher Scientific, Waltham, MA, USA) and surface antigens were subsequently assessed with the fluorochrome-conjugated antibodies listed in Table S3. For detection of intracellular proteins, surface antigen-labeled cells were fixed and permeabilized with the eBioscience Foxp3/Transcription Factor Staining Buffer Set (Thermo Fisher Scientific) and incubated with the fluorochrome-labeled antibodies listed in Table S3. All macaque samples were evaluated in a single experiment.

### Human PBMC Phenotyping by Flow Cytometry

Cryopreserved PBMCs were thawed as described previously (*78, 79, 81*), and 2-4 million cells were stained with MHC Class I multimers followed by staining for surface and then intracellular proteins. HIV-specific CD8+ T cells were identified via staining with peptide-MHC Class I multimers, either monomers that were provided by RPS and tetramerized as described (*82*) using BV421 or PE, or biotinylated pentamers (ProImmune) followed by staining with streptavidin BV421 or PE. The multimers used included MHC Class I alleles HLA-A*02, *03, *24 or B*07 plus peptides derived from Gag, Pol, Env, or Nef. PBMCs were incubated with multimer diluted in PBS for 15 minutes at room temperature (pentamer) or 20 minutes at 37°C (tetramer), followed by 20 minutes at room temperature with surface antibodies (with streptavidin for stains with pentamer) along with fixable viability dye to allow for discrimination of dead cells (Thermo Fisher) and then fixation and permeabilization using the eBioscience Foxp3/Transcription Factor Staining Buffer Set according to manufacturer’s instructions (Thermo Fisher Scientific). Antibodies used are listed in Table S3. Cytometer settings were standardized between experiments using Application settings. PBMCs from a single HIV-uninfected individual (processed and cryopreserved on the same day) were run in parallel in most experiments. Flow cytometry panels were optimized using fluorescence-minus-one controls, and gating for most markers of interest was standardized between experiments by fixing a positive gate according to the proportion of naïve (CD45RA+CCR7+CD27+) and/or effector memory CD8+ T cells (CD45RA-CCR7-CD27) from the control donor that expressed the marker. When longitudinal samples were available from a study participant, all samples were stained and run on the flow cytometer on the same experiment day. All experiments included samples from individuals from all three clinical groups. Human flow cytometry data were collected across five (Figures 1-3, 4D-H), 5A, four (Figure 4A-C), three (Figures 4D-F, H-K), or one (Figures 4C, 5B) experiment(s). Data were analyzed using FlowJo v10.1 software (Tree Star, Ashland, OR, USA).

### Proliferation by CellTrace Violet (CTV)

Thawed PBMCs in R10 media were rested overnight at 37°C and 5% CO2. Cells were then labeled with CellTrace Violet (CTV; Thermo Fisher Scientific) according to the manufacturer’s instructions. 1 million CTV-labeled cells were stimulated for five or six days (as indicated) at 37°C with 5% CO2 in R10 media containing 0.2 μg/mL HIV peptide pools comprised of peptides derived from the Gag, Nef, Env, or Pol HIV proteins (depending on the multimer specificity; supplied by the NIH AIDS Reagent Program) or individual HIV- or CMV-specific peptides (if indicated; ProImmune), or αCD3/CD28 per manufacturer’s instructions (ImmuCult human CD3/CD28 T cell activator; StemCell Technologies, Cambridge, MA). After stimulation, the cells were then stained for multimer, surface markers, and intracellular molecules, and then analyzed immediately on a flow cytometer.

### Statistical Analysis of Flow Cytometry Data

For flow cytometry data, non-parametric statistical analysis was performed using GraphPad Prism v7 (GraphPad Software, La Jolla, CA, USA) and STATA v14 (Stata Corp., College Station, TX, USA). Comparisons of parameter measurements between two separate groups of participants were assessed with Wilcoxon Rank Sum tests and between three or more separate groups were assessed with Kruskal-Wallis test followed by Dunn’s multiple comparisons test. In some cases, the same participant contributed data at multiple different timepoints in the presence and absence of ART (five individuals had samples available both pre- and post-ART initiation, and two individuals had longitudinal samples available post-ART). In these cases, linear mixed models with random intercepts were performed to control for clustering by participant, log-transforming outcome variables to satisfy model assumptions. Instances where this approach was used are highlighted in the figure legends. Differences between two subsets within individuals were assessed with paired Wilcoxon Signed Rank tests. Correlations between continuous variables were assessed with Spearman’s Rank correlations.

### Digital Droplet PCR Quantification of Cell-Associated HIV DNA

Genomic DNA was extracted from 1×10^7 PBMCs using the AllPrep DNA/RNA/miRNA Universal Kit (Qiagen, Hilden, Germany). Following nucleic acid isolation, DNA was sheared into 3kb fragments using the Covaris M220 Focused-Ultrasonicator for 10 minutes. Absolute quantification of the HIV Long Terminal Repeat (LTR) was performed, along with Ribonuclease P (RNaseP) as a cell counter, in a duplex-digital droplet PCR reaction using the Raindrop System (Raindance Technologies, Billerica, MA and Bio-Rad, Hercules, CA, USA). The Raindrop Source instrument was used to generate uniform aqueous droplets (5 picoliter) for each sample on an 8-well microfluidic chip (Raindrop Source Chip). Reactions were carried out in 50 mL volumes with 2 mL 25x Droplet Stabilizer (Raindance Technologies), 25ul 2x Taqman Genotyping Master Mix (Thermo Fisher Scientific), 900nm primers, 250nm probes and 500ng nucleic acid. Droplets were thermocycled at 95°C for 10 minutes, 45 cycles of 95°C for 15 second, 59°C for 1 minute, followed by 98°C for 10 minutes. Droplets were then placed in the deck of the Raindrop Sense instrument containing a second microfluidic chip used for single droplet fluorescence detection. Data were analyzed using the Raindrop Analyst Software on a two-dimensional histogram with FAM intensity on the X-axis and Vic intensity on the Y-axis. Cell counts were normalized to RNaseP molecules detected, with a no-template control loaded on each chip. HIV-LTR specific primers: F522-43 (5’ GCC TCA ATA AAG CTT GCC TTG A 3’; HXB2 522–543) and R626-43 (5’ GGG CGC CAC TGC TAG AGA 3’; 626–643) coupled with a FAM-BQ probe (5’ CCA GAG TCA CAC AAC AGA CGG GCA CA 3). RNase P specific primers: (5’ AGA TTT GGA CCT GCG AGC G 3’) and (5’ GAG CGG CTG TCT CCA CAA GT 3’) coupled with a VIC-MGB probe (5’ TTC TGA CCT GAA GGC TCT GCG CG 3’).

### HIV Viral Sequencing

HIV viral RNA was extracted from 140µL of plasma using the QIAmp Viral RNA Mini Kit (Qiagen). Isolated RNA was treated with TURNO DNase (Thermo Fisher Scientific) and quantified with a NanoDrop Spectrophotometer ND-1000 (Thermo Fisher Scientific). 8 µL viral RNA were reverse-transcribed into cDNA with random primers and the SuperScript III First-Strand Synthesis System (Thermo Fisher Scientific) according to the manufacturer’s instructions. Amplification of the HIV gag gene was performed using flanking primers; (5’ AAA TCT CTA GCA GTG GCG CC 3’; HXB2 623-642) and (5’ TGT TGG CTC TGG TCT GCT CT 3’; HXB2 2157-2138). PCR was performed with a Phusion High-Fidelity PCR Kit (New England Biolabs, Ipswich, MA, USA) in 20µL reactions mixed with 2µL cDNA and 10µM primers. Thermal cycling was carried out at 98°C for 30 seconds, 35 cycles of 98°C for 5 seconds, 64°C for 15 seconds, 72°C for 1 minute and finally 72°C for 10 minutes. Amplicons were purified using the QIAquick PCR purification kit (Qiagen) and Sanger sequenced by MCLABS (South San Francisco, CA, USA). Sequences were analyzed with CodonCode Aligner using a base call threshold set to identify HIV escape variants of targeted epitopes. SLYNTVATL Sequencing Primer: (5’ ACT AGC GGA GGC TAG AA 3’).

### CRISPR-Cas9 Knock Out of TCF7 in Primary Human T cells

Thawed PBMCs in R10 media were rested overnight (incubation conditions for resting or stimulating cells were 37°C and 5% CO2) and then prepared for nucleofection based on protocols described previously (*83*). Briefly, Cas9 ribonuclear proteins (RNPs) prepared by first incubating 160uM tracrRNA with 160uM *TCF7*-targeting or non-targeting (scramble) gRNA (Dharmacon, Lafayette, CO, USA) for 30 min at 37°C, then incubating this 80uM RNA product with an equal volume of 40uM Cas9 (QB3 MacroLab, University of California, Berkeley, Berkeley, CA, USA; see Table S5 for gRNA sequences). Up to 20 million PBMCs were pelleted and re-suspended in a final volume of 92uL electroporation buffer P2 (Lonza, Basel, Switzerland), combined with 8uL of the Cas9 RNPs, nucleofected using X-unit large-format cuvettes in a Lonza 4D nucleofector (pulse code EO-100), re-suspended in 1mL R10 buffer, rested for at least 15 min, and then brought to 1-2 million PBMCs/mL in R10 buffer plus 500pg/mL rhIL-7 (R&D Systems, Minneapolis, MN, USA) and rested again for four days. Cells were then harvested, an aliquot was stained for flow cytometry, and the remaining cells were labeled with CTV as described above. To provide fresh feeder cells to the cell culture, autologous (non-CTV-labeled) PBMCs thawed the day prior were depleted of CD8+ T cells (using the flow-through from the StemCell EasySep Human CD8 Positive Selection kit), rested at 37°C and 5% CO2, and then combined in a 1:1 ratio with the edited CTV-labeled PBMCs with or without αCD3/CD28 stimulatory cocktail and incubated for five days. Cells were then harvested and stained for flow cytometry. The proportion of proliferating cells (%CTV^lo^) was calculated as a proportion of total edited cells (which were gated based on positive staining for CD8α and CTV; unedited feeder cells were negative for these markers).

### CRISPR-Cas9 Generation of HIV-specific TCR-T Cells with *TCF7* Knock-in

HIV-specific TCR-T cells with *TCF7* knock-in were generated using a previously described protocol (*58*), summarized briefly and with the following modifications. Fresh primary human T cells were isolated and cultured with anti-human CD3/CD28 beads (Dynabeads, ThermoFisher), 500U/mL IL-2 (Proleukin, Prometheus Laboratories, San Diego, CA), 5ng/mL each of IL-7 (R&D Systems) and IL-15 (R&D Systems) for 48-56 hours, and then beads were magnetically separated from the T cells. In parallel, double-stranded homology directed repair (HDR) DNA templates that contained the following sequences were produced: (1) Genes encoding TCRα and TCRβ proteins derived from sequencing of an endogenous HIV-specific CD8+ T cell known to recognize the Gag/SL9 peptide presented in the context of HLA-A*02 (using a protocol previously described (*84*)); plus (2) a sequence for either the full length human *TCF7* versus truncated *NGFR* gene (see Table S5 for all HDR DNA template sequences). HDR template was combined with RNPs as reported previously (*85*) and as follows: 500ng HDR + 0.625µL crRNA ([stock] = 160µM) + 0.625µL tracrRNA ([stock] = 160µM) + 1µL poly-L-glutamic acid ([stock] = 125mg/mL) + 1.25µL Cas9 ([stock] = 40µM). 750,000 de-beaded T cells were re-suspended in 20µL Lonza electroporation buffer P3 and combined with the RNPs + HDR template and electroporated using a Lonza 4D nucleofector (pulse code EH-115). Cells were rescued in media (Lonza X-Vivo 15 media with 5% FCS, NAC, 2-mercaptoethanol) without cytokines for 10 min at 37°C and then cultured in media (X-Vivo 15 media with 5% FCS, NAC, 2-mercaptoethanol) plus 500 U/ml IL-2 for seven days with exchange of media and IL-2 every two days. After seven days, an aliquot of the cells was harvested and stained for flow cytometry. Edited T cells were CTV labeled and cultured at an effector:target ratio of 1:1 (effector numbers based on the percent of live cells that were HLA-A*02:SL9 pentamer+) with an HLA-A*02 cell line as target/antigen presenting cells (T2 cells; ATCC, Old Town Manassa, VA, USA) coated with 1µM SL9 peptide (ProImmune). Cells were co-incubated for six days and then stained for evaluation by flow cytometry.

## Supplementary Materials

**Figure S1.**
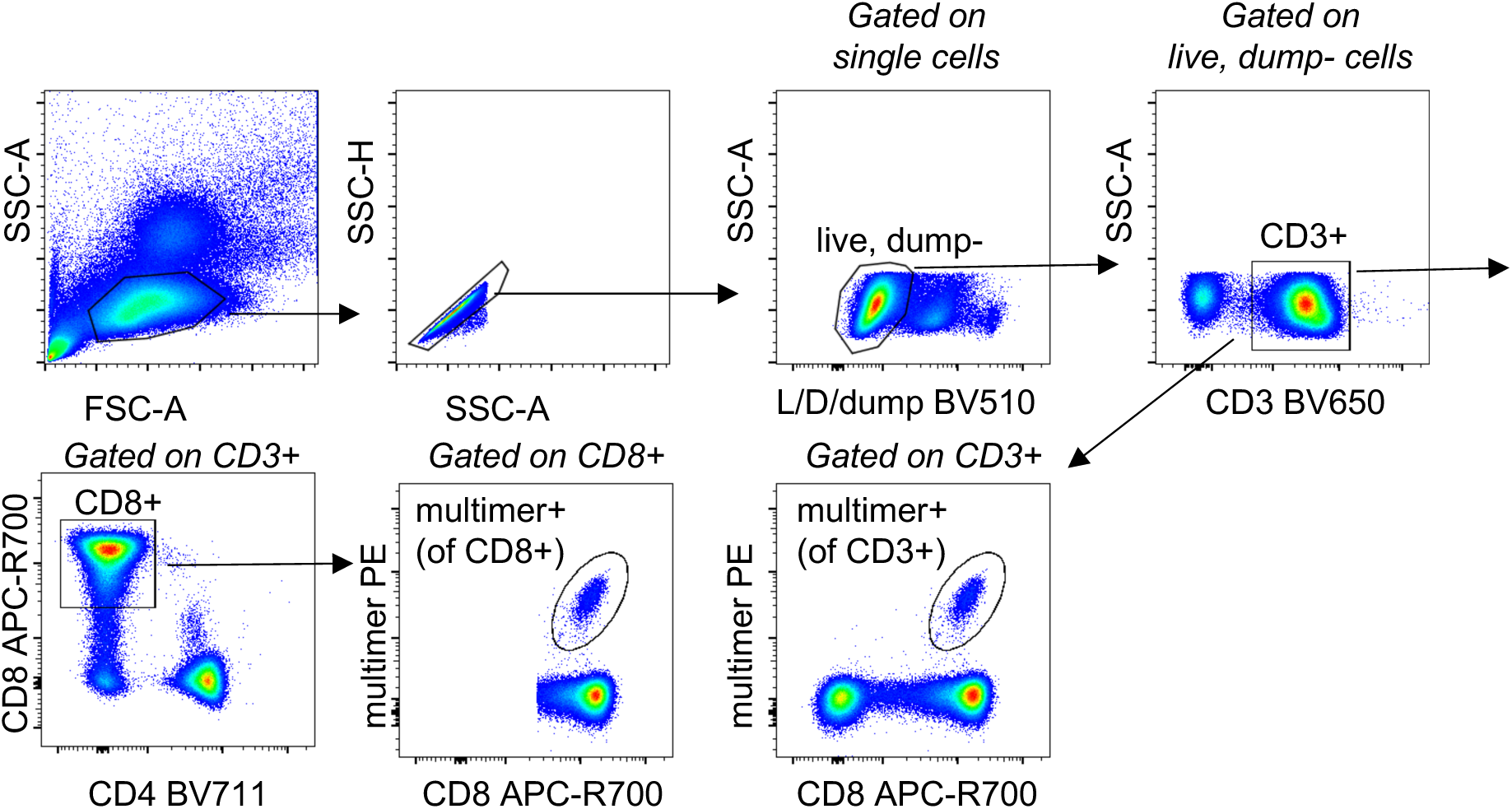
MHC Class I multimer+ HIV-specific CD8+ T cell gating strategy. Dump gate includes staining for CD14, CD19 and TCR-γδ.

**Figure S2.**
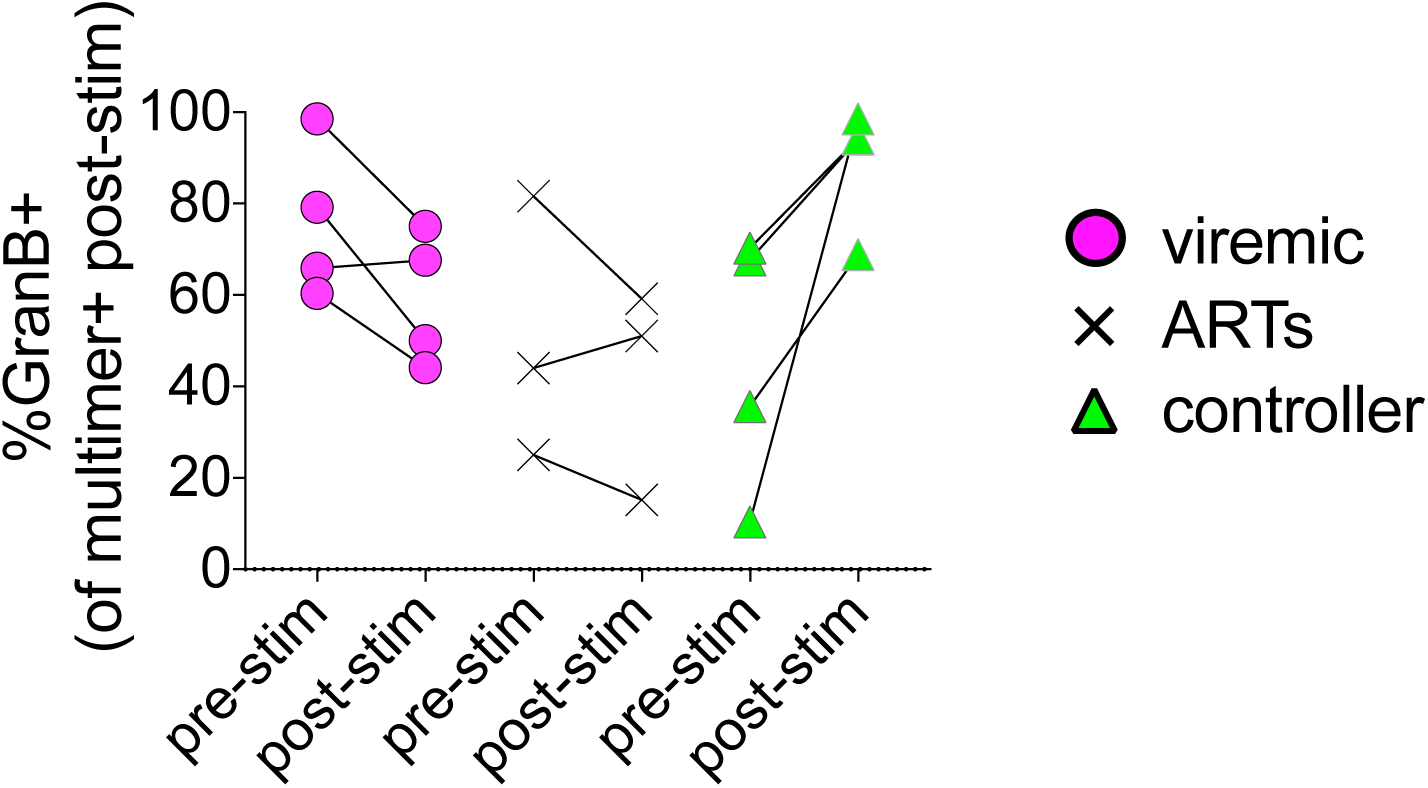
Change in Granzyme B expression after six-day *in vitro* peptide stimulation of multimer+ HIV-specific CD8+ T cells. Percentage of Granzyme B+ multimer+ cells prior to and after stimulation.

**Figure S3.**
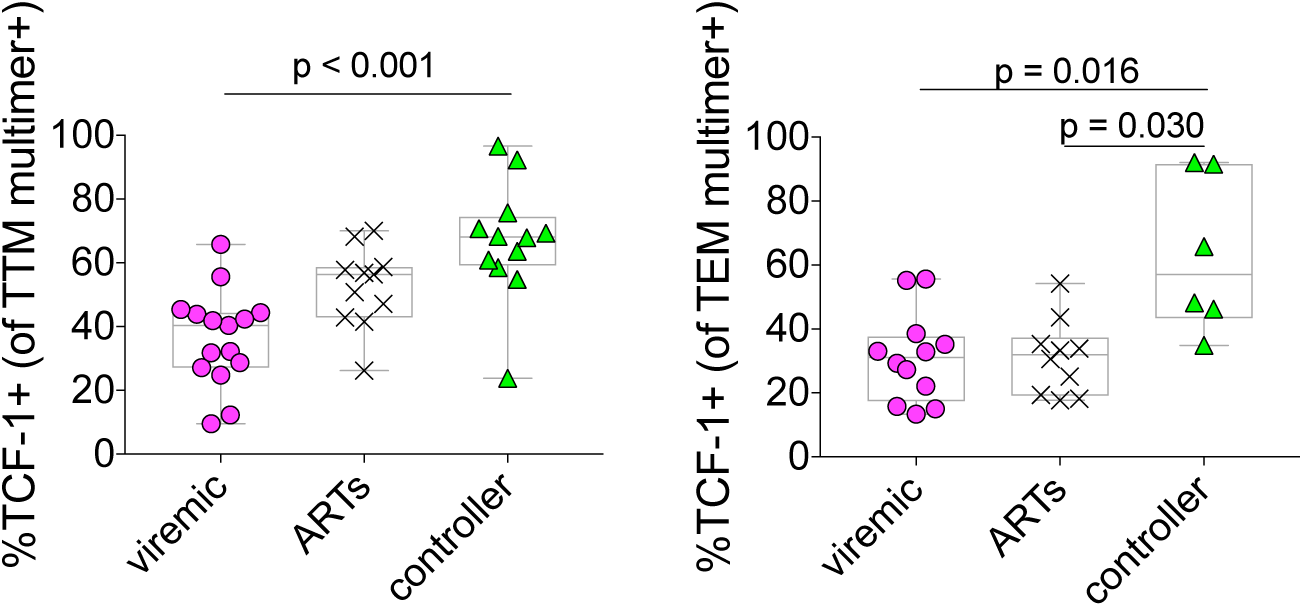
TCF-1 expression amongst TTM and TEM multimer+ HIV-specific CD8+ T cells. Statistical testing: Linear mixed effects models to account for clustering within participants.

**Figure S4.**
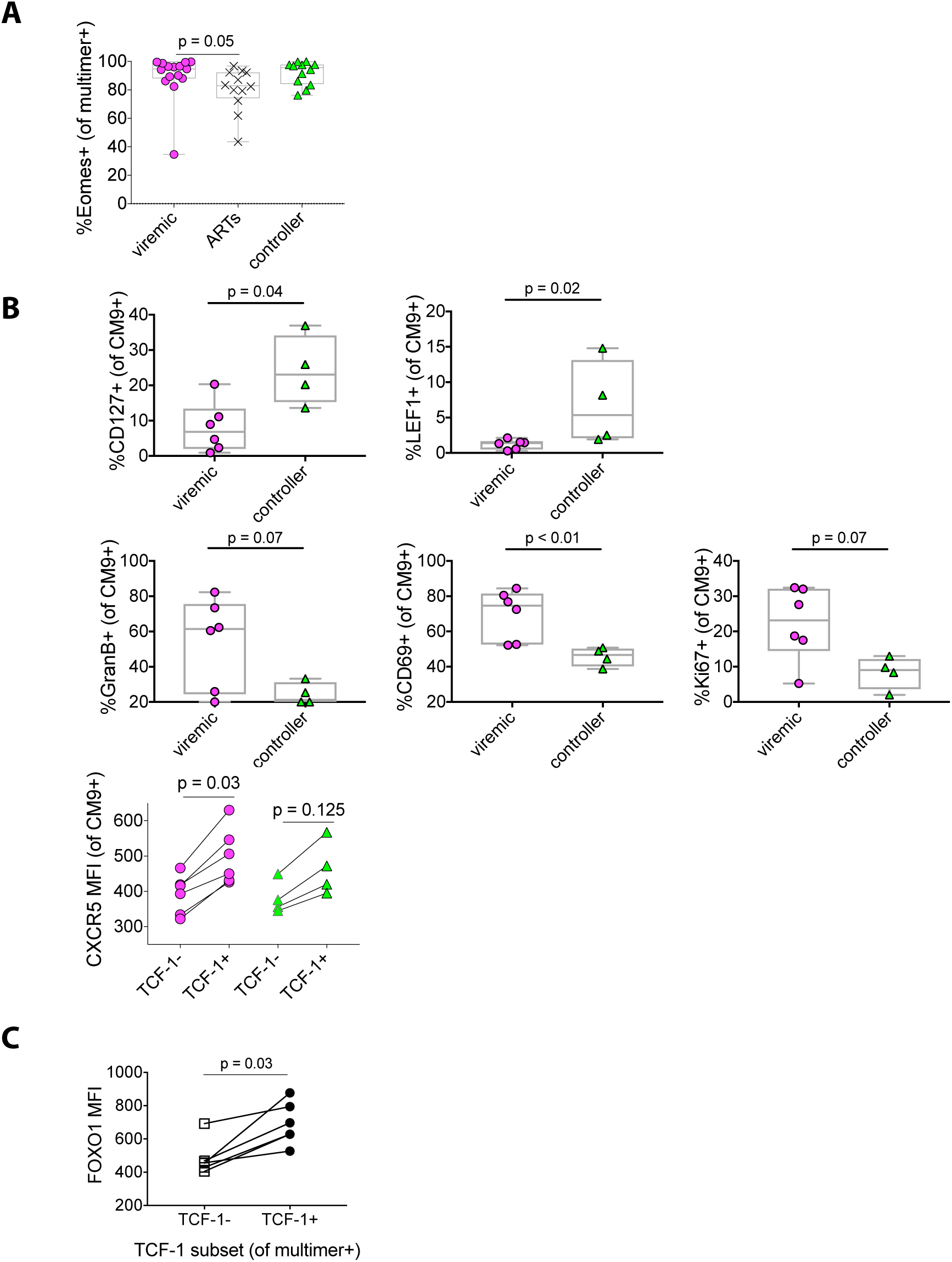
Phenotypes of multimer+ HIV-specific and SIV-specific CD8+ T cells. (**A**) Expression of Eomesodermin (Eomes) in HIV-specific CD8+ T cells. (**B**) Phenotype of SIV-specific CD8+ T cells from viremic and controller animals. (**C**) Expression of FOXO1 in TCF-1+ and TCF-1-subsets of multimer+ HIV-specific CD8+ T cells. Statistical testing: Linear mixed effects models to account for clustering within participants (A), Wilcoxon Rank Sum (B), Wilcoxon Signed Rank (paired data in B; C).

**Figure S5.**
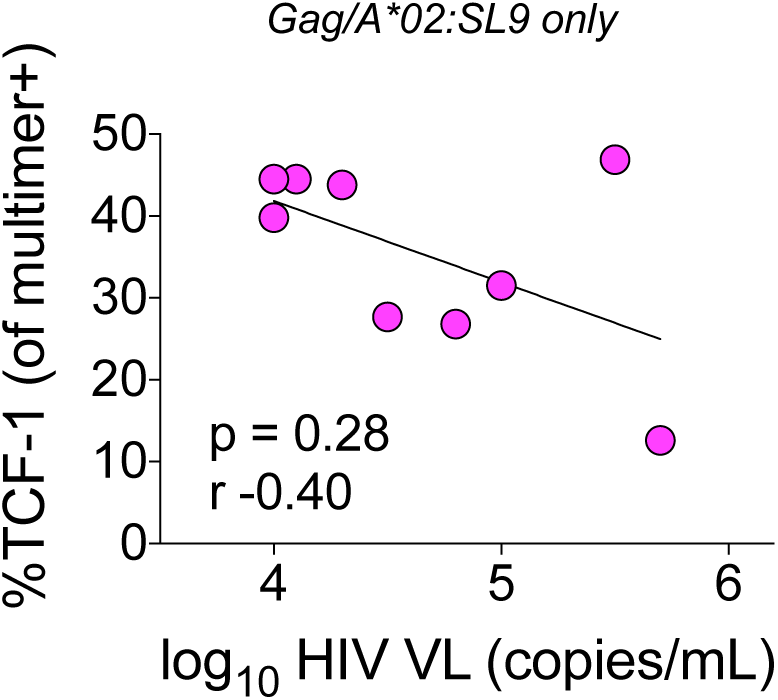
Correlation between TCF-1 expression in multimer+ HIV-specific CD8+ T cells and viral load amongst individuals with documented viral CD8+ T cell escape variants. Statistical testing: Spearman Correlation.

**Figure S6.**
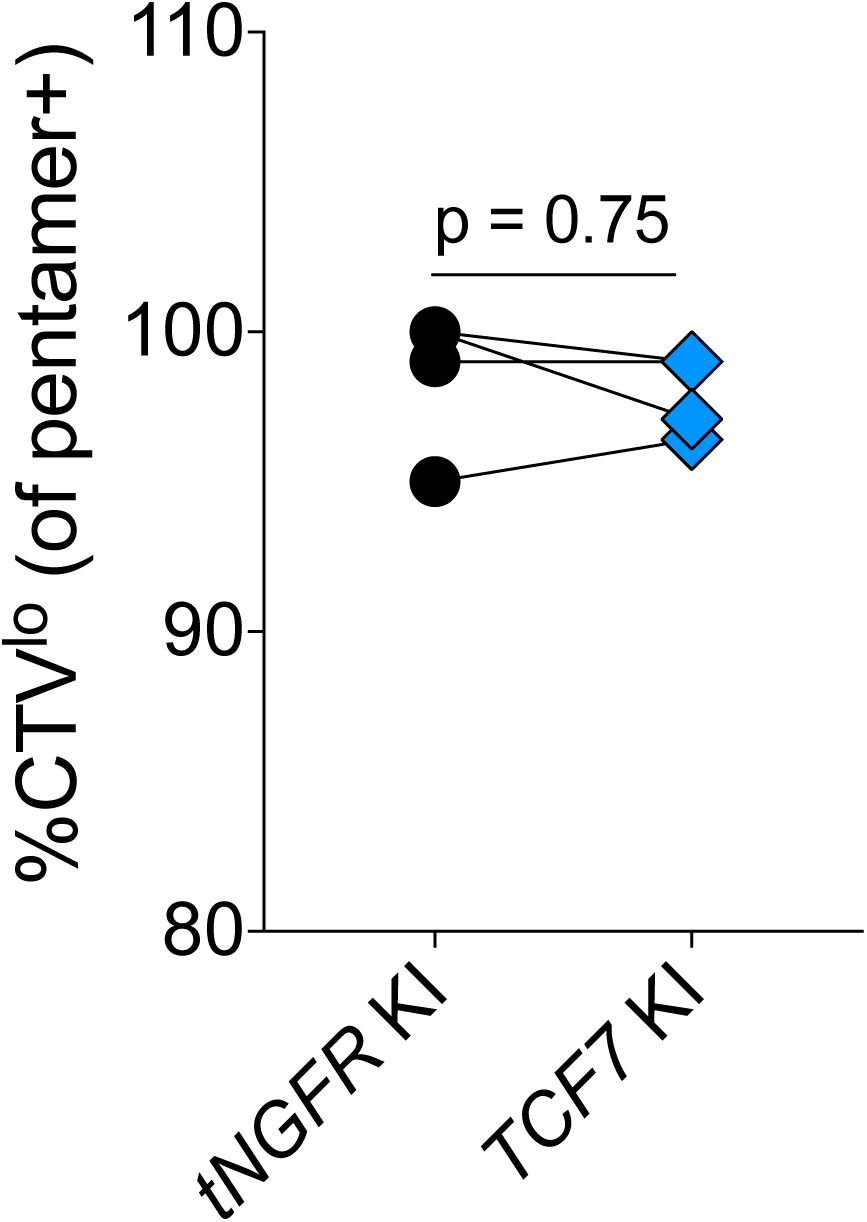
Proliferation of HIV-specific TCR-T cells. Proportion of divided cells with *tNGFR* versus *TCF7* overexpression after 6-day stimulation with cognate peptide-loaded antigen presenting cells. Statistical testing: Wilcoxon Signed Rank test.

**Table S1.**
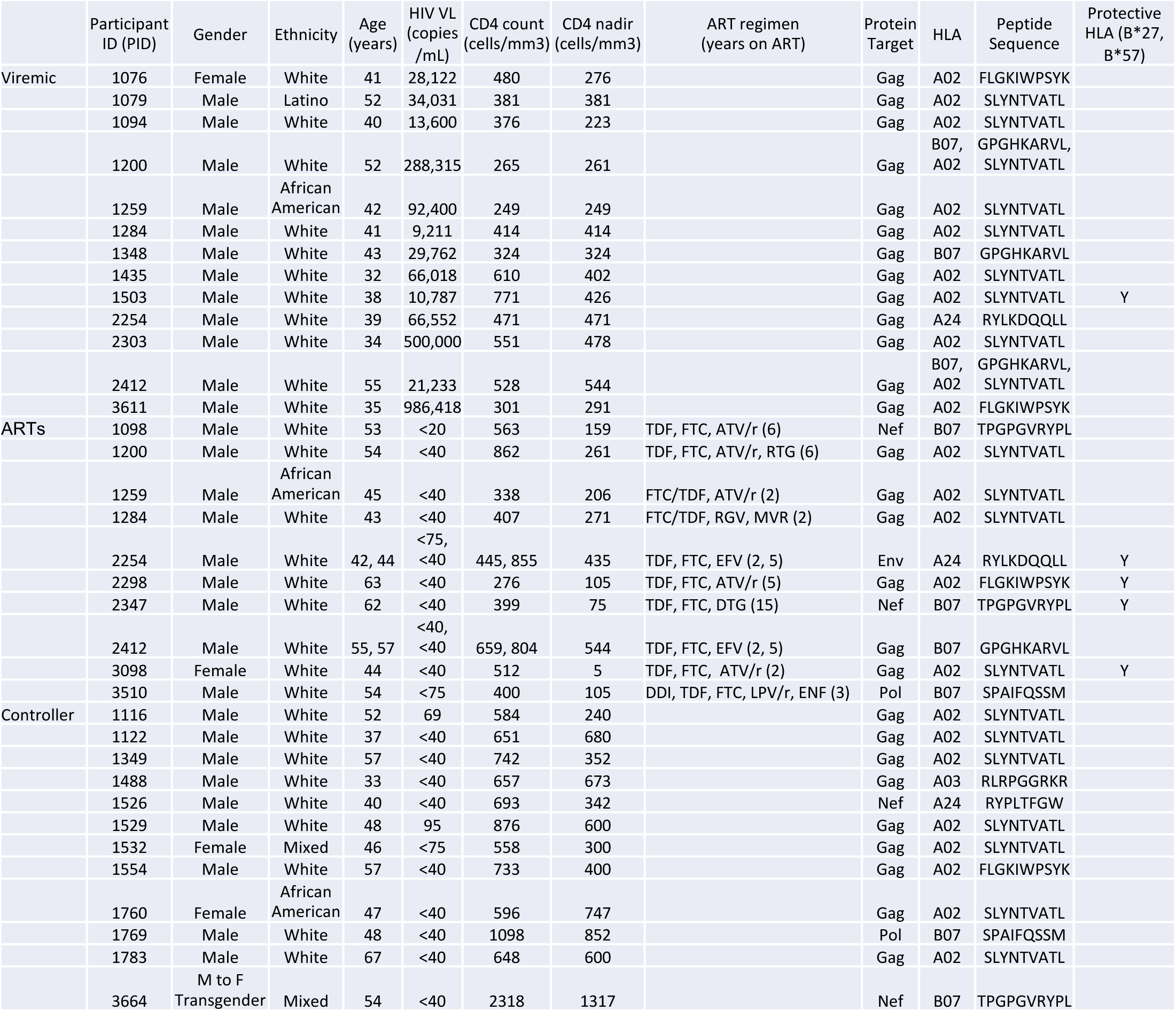
SCOPE participant clinical information. PID, participant ID. TDF, tenofovir disoproxil. FTC, emtricitabine. ATV/r, atazanavir/ritonavir. RGV, raltegravir. MVR, maraviroc. EFV, efavirenz. DTG, dolutegravir. DDI, diadenosine. ENF, enfuviritide.

**Table S2.** SIV-infected rhesus macaque clinical information.

|  | Animal ID | SIV Virus | CD4 count<br>(cells/mm3) | SIV VL<br>(copies/mL) |
| --- | --- | --- | --- | --- |
| Viremic | CL4C | mac239 | 47 | 470000 |
|  | CL86 | mac239 | 91 | 900000 |
|  | 591 | smE543 | 186 | 251000 |
|  | 764 | smE543 | 134 | 222000 |
|  | 766 | smE543 | 927 | 50000 |
|  | 828 | smE543 | 264 | 330000 |
| Controller | F98 | mac239 | 560 | 15 |
|  | 863 | smE660 | 1245 | 370 |
|  | 867 | smE660 | 634 | 15 |
|  | 871 | smE660 | 513 | 15 |

**Table S3.**
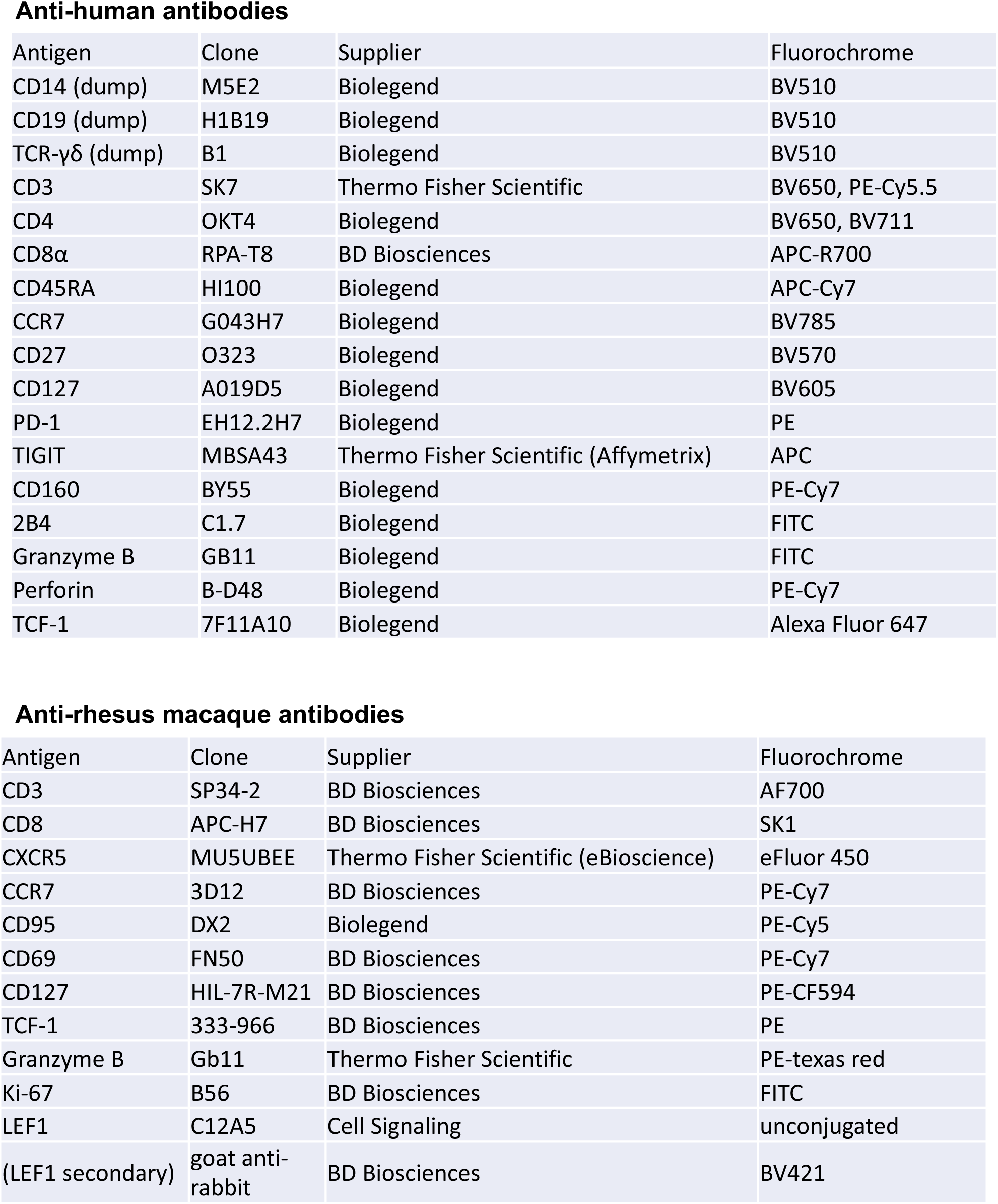
Antibodies and multimers used for flow cytometry staining.

Anti-human antibodies
| Antigen | Clone | Supplier | Fluorochrome |
| --- | --- | --- | --- |
| CD14 (dump) | M5E2 | Biologend | BV510 |
| CD19 (dump) | H1B19 | Biologend | BV510 |
| TCR-$\gamma\delta$ (dump) | B1 | Biologend | BV510 |
| CD3 | SK7 | Thermo Fisher Scientific | BV650, PE-Cy5.5 |
| CD4 | OKT4 | Biologend | BV650, BV711 |
| CD8α | RPA-T8 | BD Biosciences | APC-R700 |
| CD45RA | HI100 | Biologend | APC-Cy7 |
| CCR7 | G043H7 | Biologend | BV785 |
| CD27 | O323 | Biologend | BV570 |
| CD127 | A019D5 | Biologend | BV605 |
| PD-1 | EH12.2H7 | Biologend | PE |
| TIGIT | MBSA43 | Thermo Fisher Scientific (Affymetrix) | APC |
| CD160 | BY55 | Biologend | PE-Cy7 |
| 2B4 | C1.7 | Biologend | FITC |
| Granzyme B | GB11 | Biologend | FITC |
| Perforin | B-D48 | Biologend | PE-Cy7 |
| TCF-1 | 7F11A10 | Biologend | Alexa Fluor 647 |

**Table S3. Antibodies and multimers used for flow cytometry staining.**
| Antigen | Clone | Supplier | Fluorochrome |
| --- | --- | --- | --- |
| CD3 | SP34-2 | BD Biosciences | AF700 |
| CD8 | APC-H7 | BD Biosciences | SK1 |
| CXCR5 | MU5UBEE | Thermo Fisher Scientific (eBioscience) | eFluor 450 |
| CCR7 | 3D12 | BD Biosciences | PE-Cy7 |
| CD95 | DX2 | Biologend | PE-Cy5 |
| CD69 | FN50 | BD Biosciences | PE-Cy7 |
| CD127 | HIL-7R-M21 | BD Biosciences | PE-CF594 |
| TCF-1 | 333-966 | BD Biosciences | PE |
| Granzyme B | Gb11 | Thermo Fisher Scientific | PE-texas red |
| Ki-67 | B56 | BD Biosciences | FITC |
| LEF1 | C12A5 | Cell Signaling | unconjugated |
| (LEF1 secondary) | goat anti-rabbit | BD Biosciences | BV421 |

**Table S4.** HIV peptide sequence variants of wildtype HLA-A*02:Gag-SL9 9mer.

| Participant ID | Major peptide sequence | Secondary detected peptide sequence |
| --- | --- | --- |
| 1079 | SLYNTIAVL | SLYNTIATL |
| 1200 | SLYNTVAVL | SLYNTIAVL |
| 1259 | SLYNTIAVL | SLYNTVAVL / SLYNTVATL (WT) |
| 1284 | SLYNTVAVL |  |
| 1435 | SLYNTIAVL | SLYNTVAVL |
| 1503 | SLYNTIAVL |  |
| 1904 | SLYNTIAVL |  |

**Table S5.**
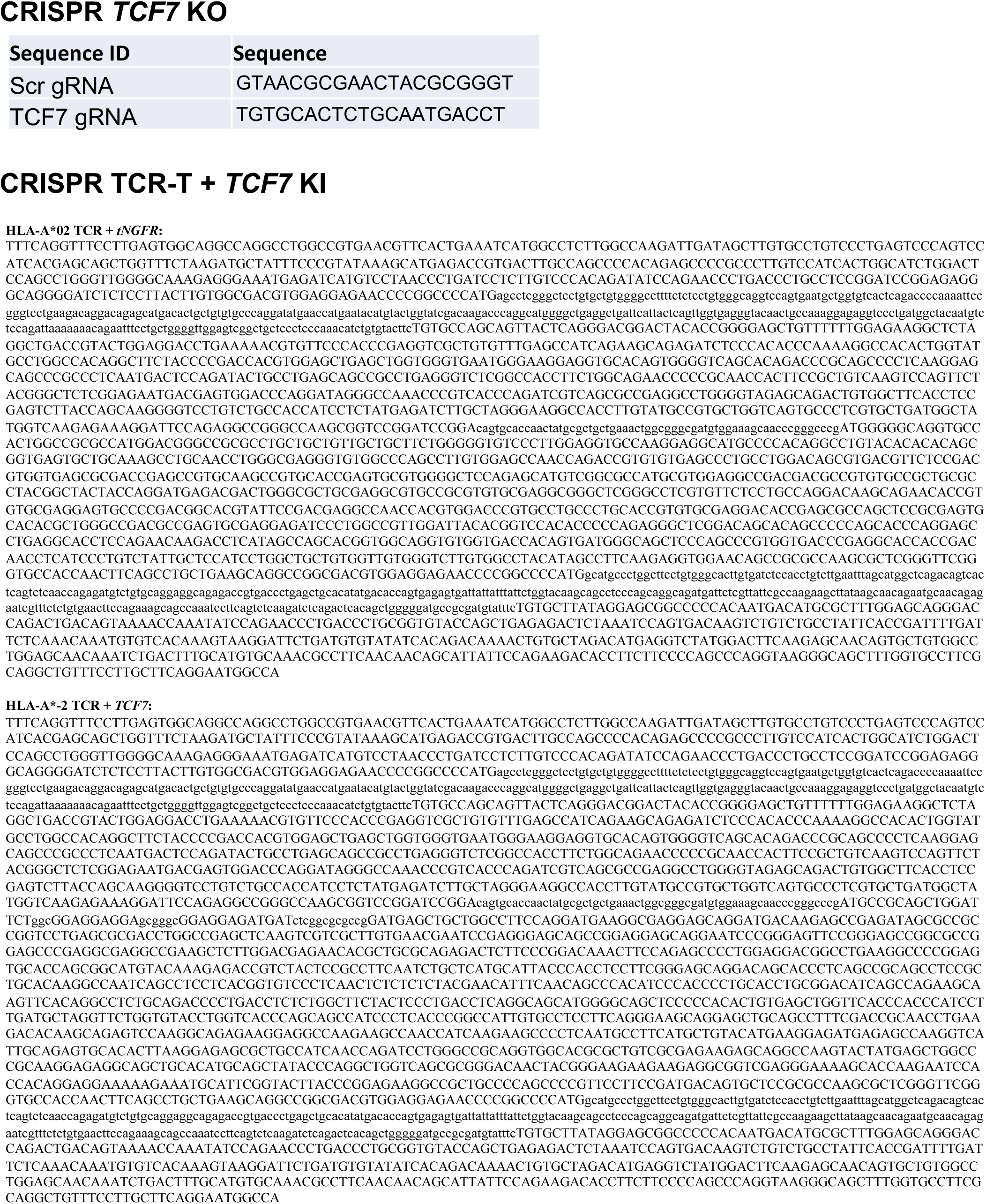
Guide RNA and HDR template sequences for CRISPR-Cas9 knock-out and knock-in experiments.

## Acknowledgements

We sincerely thank all study participants. We thank Michelle Hough and Maya Ball-Burak for assistance with accessing SCOPE samples, and Devin Cavero, Ventura Mendoza, and Erin Isaza with reagent preparation.

## Funding

This work was supported by the National Institutes of Health: 1K23AI134327 and 5T32AI060530 (R.L.R), The Delaney AIDS Research Enterprise to find a cure (DARE; 5U19A1096109, A127966; S.G.D., P.W.H., J.M.M., R.L.R.), 5K24AI069994 (S.G.D.), R01HD074511 (C.D.P.), R01AI110271 (P.W.H.), P50GM082250 (F.B, A.M.), and the Division of Intramural Research, National Institute of Allergy and Infectious Diseases (C.E.S., J.C.M., and J.M.B.). The SCOPE cohort is supported the UCSF/Gladstone Institute of Virology & Immunology CFAR (P30AI027763), the UCSF Clinical and Translational Research Institute Clinical Research Center (UL1TR001872), and the CFAR Network of Integrated Systems (R24AI067039). Additional support for SCOPE was provided by DARE and the amfAR Institute for HIV Cure Research (amfAR 1093t01). Options cohort support was also provided by the Bill and Melinda Gates Foundation (OPP1062806) and the Harvey V. Berneking Living Trust. C.D.T.D. received support from the Philippine Department of Science and Technology. F.B. was supported by the Care-for-Rare Foundation and the German Research Foundation (D.F.G.). A.M. holds a Career Award for Medical Scientists from the Burroughs Wellcome Fund, is an investigator at the Chan Zuckerberg Biohub, and has received funding from the Innovative Genomics Institute (IGI) and the Parker Institute for Cancer Immunotherapy (PICI).

## Author contributions

R.L.R designed the study and performed all flow cytometry and cell sorting experiments with assistance from C.D.T.D., W.C., C.S., and L.W. R.P.S. provided MHC Class I monomers. CRISPR KO experiments were performed by J.H. and R.L.R. and CRISPR KI experiments were performed by R.L.R., F.B., and L.W. with input from A.M and T.R. HIV-specific TCR sequences were provided by R.M-N., D.C.D., and R.A.K. Viral sequencing and reservoir measurements were performed by K.R. and analysis was overseen by S.K.P. SCOPE and Options samples were kindly provided by S.G.D., J.N.M., R.M.H., and C.D.P. with cohort data management support from R.H. and M.K. The manuscript was written by R.L.R. P.W.H., J.M.M., S.G.D., and R.P.S. provided scientific guidance throughout.

## Competing interests

The authors declare competing financial interests: T.L.R. and A.M. are co-founders of Arsenal Biosciences. T.L.R. served as the CSO of Arsenal Biosciences from March to December 2019. A.M. is a co-founder of Spotlight Therapeutics. A.M. serves as on the scientific advisory board of PACT Pharma, is an advisor to Trizell, and was a former advisor to Juno Therapeutics. The Marson Laboratory has received sponsored research support from Juno Therapeutics, Epinomics, Sanofi and a gift from Gilead.

## Data and materials availability

All raw flow cytometry files will be provided upon request.

## References and Notes

1. L. Trautmann, L. Janbazian, N. Chomont, E. A. Said, S. Gimmig, B. Bessette, M.-R. Boulassel, E. Delwart, H. Sepulveda, R. S. Balderas, J.-P. Routy, E. K. Haddad, R.-P. Sekaly. Upregulation of PD-1 expression on HIV-specific CD8+ T cells leads to reversible immune dysfunction., Nat. Med. 12, 1198–1202 (2006).

2. C. L. Day, D. E. Kaufmann, P. Kiepiela, J. A. Brown, E. S. Moodley, S. Reddy, E. W. Mackey, J. D. Miller, A. J. Leslie, C. DePierres, Z. Mncube, J. Duraiswamy, B. Zhu, Q. Eichbaum, M. Altfeld, J. E. Wherry, H. M. Coovadia, P. J. Goulder, P. Klenerman, R. Ahmed, G. J. Freeman, B. D. Walker. PD-1 expression on HIV-specific T cells is associated with T-cell exhaustion and disease progression. Nature. 443, 350–354 (2006).

3. S. Kostense, G. Ogg, E. Manting, G. Gillespie, J. Joling, K. Vandenberghe, E. Veenhof, D. van Baarle, S. Jurriaans, M. Klein, F. Miedema, High viral burden in the presence of major HIV-specific CD8(+) T cell expansions: evidence for impaired CTL effector function. Eur. J. Immunol. 31, 677–686 (2001).

4. S. G. Deeks, B. D. Walker, Human Immunodeficiency Virus Controllers: Mechanisms of Durable Virus Control in the Absence of Antiretroviral Therapy. Immunity. 27, 406–416 (2007).

5. P. Borrow, H. Lewicki, B. Hahn, G. aw, M. Oldstone, Virus-specific CD8+ cytotoxic T-lymphocyte activity associated with control of viremia in primary human immunodeficiency virus type 1 infection. J. Virol. 68, 6103–6110 (1994).

6. J. Cao, J. McNevin, U. Malhotra, J. M. McElrath, Evolution of CD8+ T cell immunity and viral escape following acute HIV-1 infection. J. Immunol. 171, 3837–3846 (2003).

7. C. Riou, W. A. Burgers, K. Mlisana, R. A. Koup, M. Roederer, S. S. Karim, C. Williamson, C. M. Gray, Differential impact of magnitude, polyfunctional capacity, and specificity of HIV-specific CD8+ T cell responses on HIV set point. J. Virol. 88, 1819–1824 (2014).

8. Z. M. Ndhlovu, P. Kamya, N. Mewalal, H. N. Kløverpris, T. Nkosi, K. Pretorius, F. Laher, F. Ogunshola, D. Chopera, K. Shekhar, M. Ghebremichael, N. Ismail, A. Moodley, A. Malik, A. Leslie, P. J. Goulder, S. Buus, A. Chakraborty, K. Dong, T. Ndung’u, B. D. Walker. Magnitude and Kinetics of CD8(+) T Cell Activation during Hyperacute HIV Infection Impact Viral Set Point. Immunity. 43, 591–604 (2015).

9. H. Takata, S. Buranapraditkun, C. Kessing, J. L. Fletcher, R. Muir, V. Tardif, P. Cartwright, C. Vandergeeten, W. Bakeman, C. N. Nichols, S. Pinyakorn, P. Hansasuta, E. Kroon, T. Chalermchai, R. O’Connell, J. Kim, N. Phanuphak, M. L. Robb, N. L. Michael, N. Chomont, E. K. Haddad, J. Ananworanich, L. Trautmann, R. and the RV254/SEARCH010 and RV304/SEARCH013 Study Groups. Delayed differentiation of potent effector CD8(+) T cells reducing viremia and reservoir seeding in acute HIV infection. Sci. Transl. Med. 9, eaag1809 (2017).

10. G. Mylvaganam, A. G. Yanez, M. Maus, B. D. Walker. Toward T Cell-Mediated Control or Elimination of HIV Reservoirs: Lessons From Cancer Immunology. Front. Immunol. 10, 2109 (2019).

11. J. A. Warren, G. Clutton, N. Goonetilleke. Harnessing CD8+ T Cells Under HIV Antiretroviral Therapy. Front. Immunol. 10, 291 (2019).

12. B. R. Jones, B. D. Walker. HIV-specific CD8^+^ T cells and HIV eradication. J. Clin. Invest. 126, 455–463 (2016).

13. L. M. McLane, M. S. Abdel-Hakeem, J. E. Wherry. CD8 T Cell Exhaustion During Chronic Viral Infection and Cancer. Annu. Rev. Immunol. 37, 457–495 (2019).

14. International HIV Controllers Study, F. Pereyra, X. Jia, P. J. McLaren, P. I. de Bakker, B. D. Walker, S. Ripke, C. J. Brumme, S. L. Pulit, M. Carrington, C. M. Kadie, J. M. Carlson, D. Heckerman, R. R. Graham, R. M. Plenge, S. G. Deeks, L. Gianniny, G. Crawford, J. Sullivan, E. Gonzalez, L. Davies, A. Camargo, J. M. Moore, N. Beattie, S. Gupta, A. Crenshaw, N. P. Burtt, C. Guiducci, N. Gupta, X. Gao, Y. Qi, Y. Yuki, A. Piechocka-Trocha, E. Cutrell, R. Rosenberg, K. L. Moss, P. Lemay, J. O’Leary, T. Schaefer, P. Verma, I. Toth, B. Block, B. Baker, A. Rothchild, J. Lian, J. Proudfoot, D. L. Alvino, S. Vine, M. M. Addo, T. M. Allen, M. Altfeld, M. R. Henn, S. Gall, H. Streeck, D. W. Haas, D. R. Kuritzkes, G. K. Robbins, R. W. Shafer, R. M. Gulick, C. ikuma, R. Haubrich, S. Riddler, P. E. Sax, E. S. Daar, H. J. Ribaudo, B. Agan, S. Agarwal, R. L. Ahern, B. L. Allen, S. Altidor, E. L. Altschuler, S. Ambardar, K. Anastos, B. Anderson, V. Anderson, U. Andrady, D. Antoniskis, D. Bangsberg, D. Barbaro, W. Barrie, J. Bartczak, S. Barton, P. Basden, N. Basgoz, S. Bazner, N. C. Bellos, A. M. Benson, J. Berger, N. F. Bernard, A. M. Bernard, C. Birch, S. J. Bodner, R. K. Bolan, E. T. Boudreaux, M. Bradley, J. F. Braun, J. E. Brndjar, S. J. Brown, K. Brown, S. T. Brown, J. Burack, L. M. Bush, V. Cafaro, O. Campbell, J. Campbell, R. H. Carlson, K. J. Carmichael, K. K. Casey, C. Cavacuiti, G. Celestin, S. T. Chambers, N. Chez, L. M. Chirch, P. J. Cimoch, D. Cohen, L. E. Cohn, B. Conway, D. A. Cooper, B. Cornelson, D. T. Cox, M. V. Cristofano, G. Cuchural, J. L. Czartoski, J. hman, J. S. Daly, B. T. Davis, K. Davis, S. vod, E. Dejesus, C. A. Dietz, E. Dunham, M. E. Dunn, T. B. Ellerin, J. J. Eron, J. J. Fangman, C. E. Farel, H. Ferlazzo, S. Fidler, A. Fleenor-Ford, R. Frankel, K. A. Freedberg, N. K. French, J. D. Fuchs, J. D. Fuller, J. Gaberman, J. E. Gallant, R. T. Gandhi, E. Garcia, D. Garmon, J. C. Gathe, C. R. Gaultier, W. Gebre, F. D. Gilman, I. Gilson, P. A. Goepfert, M. S. Gottlieb, C. Goulston, R. K. Groger, D. T. Gurley, S. Haber, R. Hardwicke, D. W. Hardy, R. P. Harrigan, T. N. Hawkins, S. Heath, F. M. Hecht, K. W. Henry, M. Hladek, R. P. Hoffman, J. M. Horton, R. K. Hsu, G. D. Huhn, P. Hunt, M. J. Hupert, M. L. Illeman, H. Jaeger, R. M. Jellinger, M. John, J. A. Johnson, K. L. Johnson, H. Johnson, K. Johnson, J. Joly, W. C. Jordan, C. A. Kauffman, H. Khanlou, R. K. Killian, A. Y. Kim, D. D. Kim, C. A. Kinder, J. T. Kirchner, L. Kogelman, E. Kojic, T. P. Korthuis, W. Kurisu, D. S. Kwon, M. LaMar, H. Lampiris, M. Lanzafame, M. M. Lederman, D. M. Lee, J. M. Lee, M. J. Lee, E. T. Lee, J. Lemoine, J. A. Levy, J. M. Llibre, M. A. Liguori, S. J. Little, A. Y. Liu, A. J. Lopez, M. R. Loutfy, D. Loy, D. Y. Mohammed, A. Man, M. K. Mansour, V. C. Marconi, M. Markowitz, R. Marques, J. N. Martin, H. L. Martin, K. Mayer, J. M. McElrath, T. A. McGhee, B. H. McGovern, K. McGowan, D. McIntyre, G. X. Mcleod, P. Menezes, G. Mesa, C. E. Metroka, D. Meyer-Olson, A. O. Miller, K. Montgomery, K. C. Mounzer, E. H. Nagami, I. Nagin, R. G. Nahass, M. O. Nelson, C. Nielsen, D. L. Norene, D. H. O’Connor, B. O. Ojikutu, J. Okulicz, O. O. Oladehin, E. C. Oldfield, S. A. Olender, M. Ostrowski, W. F. Owen, E. Pae, J. Parsonnet, A. M. Pavlatos, A. M. Perlmutter, M. N. Pierce, J. M. Pincus, L. Pisani, L. Price, L. Proia, R. C. Prokesch, H. Pujet, M. Ramgopal, A. Rathod, M. Rausch, J. Ravishankar, F. S. Rhame, C. Richards, D. D. Richman, B. Rodes, M. Rodriguez, R. C. Rose, E. S. Rosenberg, D. Rosenthal, P. E. Ross, D. S. Rubin, E. Rumbaugh, L. Saenz, M. R. Salvaggio, W. C. Sanchez, V. njana, S. Santiago, W. Schmidt, H. Schuitemaker, P. stak, P. Shalit, W. Shay, V. N. Shirvani, V. I. Silebi, J. zemore, P. R. Skolnik, M. Sokol-Anderson, J. sman, P. Stabile, J. T. Stapleton, S. Starrett, F. Stein, H.-J. Stellbrink, L. F. Sterman, V. E. Stone, D. R. Stone, G. Tambussi, R. A. Taplitz, E. M. Tedaldi, A. Telenti, W. Theisen, R. Torres, L. Tosiello, C. Tremblay, M. A. Tribble, P. D. Trinh, A. Tsao, P. Ueda, A. Vaccaro, E. Valadas, T. J. Vanig, I. Vecino, V. M. Vega, W. Veikley, B. H. Wade, C. Walworth, C. Wanidworanun, D. J. Ward, D. A. Warner, R. D. Weber, D. Webster, S. Weis, D. A. Wheeler, D. J. White, E. Wilkins, A. Winston, C. G. Wlodaver, A. Wout, D. P. Wright, O. O. Yang, D. L. Yurdin, B. W. Zabukovic, K. C. Zachary, B. Zeeman, M. Zhao. The major genetic determinants of HIV-1 control affect HLA class I peptide presentation. Science. 330, 1551–1557 (2010).

15. C. Ekenberg, M.-H. Tang, A. G. Zucco, D. D. Murray, C. MacPherson, X. Hu, B. T. Sherman, M. H. Losso, R. Wood, R. Paredes, J.-M. Molina, M. Helleberg, N. Jina, C. M. Kityo, E. Florence, M. N. Polizzotto, J. D. Neaton, C. H. Lane, J. D. Lundgren, INSIGHT START Study Group. Association Between Single-Nucleotide Polymorphisms in HLA Alleles and Human Immunodeficiency Virus Type 1 Viral Load in Demographically Diverse, Antiretroviral Therapy–Naive Participants From the Strategic Timing of AntiRetroviral Treatment Trial. J. Infect. Dis. 220, 1325–1334 (2019).

16. P. J. McLaren, C. Coulonges, I. Bartha, T. L. Lenz, A. J. Deutsch, A. Bashirova, S. Buchbinder, M. N. Carrington, A. Cossarizza, J. Dalmau, A. Luca, J. J. Goedert, D. Gurdasani, D. W. Haas, J. T. Herbeck, E. O. Johnson, G. D. Kirk, O. Lambotte, M. Luo, S. Mallal, D. van Manen, J. Martinez-Picado, L. Meyer, J. M. Miro, J. I. Mullins, N. Obel, G. Poli, M. S. Sandhu, H. Schuitemaker, P. R. Shea, I. Theodorou, B. D. Walker, A. C. Weintrob, C. A. Winkler, S. M. Wolinsky, S. Raychaudhuri, D. B. Goldstein, A. Telenti, P. I. de Bakker, J.-F. Zagury, J. Fellay. Polymorphisms of large effect explain the majority of the host genetic contribution to variation of HIV-1 virus load. P. Natl. Acad. Sci. 112, 14658–63 (2015).

17. J. Fellay, K. V. Shianna, D. Ge, S. Colombo, B. Ledergerber, M. Weale, K. Zhang, C. Gumbs, A. Castagna, A. Cossarizza, A. Cozzi-Lepri, A. Luca, P. Easterbrook, P. Francioli, S. Mallal, J. Martinez-Picado, J. M. Miro, N. Obel, J. P. Smith, J. Wyniger, P. Descombes, S. E. Antonarakis, N. L. Letvin, A. J. McMichael, B. F. Haynes, A. Telenti, D. B. Goldstein. A whole-genome association study of major determinants for host control of HIV-1. Science. 317, 944– 947 (2007).

18. A. Chowdhury, T. L. Hayes, S. E. Bosinger, B. O. Lawson, T. Vanderford, J. E. Schmitz, M. Paiardini, M. Betts, A. Chahroudi, J. D. Estes, G. Silvestri. Differential Impact of In Vivo CD8+ T Lymphocyte Depletion in Controller versus Progressor Simian Immunodeficiency Virus-Infected Macaques. J. Virol. 89, 8677–8686 (2015).

19. S. A. Migueles, A. C. Laborico, L. W. Shupert, irin Sabbaghian, R. Rabin, C. W. Hallahan, D. van Baarle, S. Kostense, F. Miedema, M. McLaughlin, L. Ehler, J. Metcalf, S. Liu, M. Connors. HIV-specific CD8+ T cell proliferation is coupled to perforin expression and is maintained in nonprogressors. Nat. Immunol. 3, 1061–1068 (2002).

20. M. R. Betts, M. C. Nason, S. M. West, S. C. Rosa, S. A. Migueles, J. Abraham, M. M. Lederman, J. M. Benito, P. A. Goepfert, M. Connors, M. Roederer, R. A. Koup. HIV nonprogressors preferentially maintain highly functional HIV-specific CD8+ T cells. Blood. 107, 4781–4789 (2006).

21. S. A. Migueles, C. M. Osborne, C. Royce, A. A. Compton, R. P. Joshi, K. A. Weeks, J. E. Rood, A. M. Berkley, J. B. Sacha, N. A. Cogliano-Shutta, M. Lloyd, G. Roby, R. Kwan, M. McLaughlin, S. Stallings, C. Rehm, M. A. O’Shea, J. Mican, B. Z. Packard, A. Komoriya, S. Palmer, A. P. Wiegand, F. Maldarelli, J. M. Coffin, J. W. Mellors, C. W. Hallahan, D. A. Follman, M. Connors. Lytic granule loading of CD8+ T cells is required for HIV-infected cell elimination associated with immune control. Immunity. 29, 1009–1021 (2008).

22. S. A. Migueles, K. A. Weeks, E. Nou, A. M. Berkley, J. E. Rood, C. M. Osborne, C. W. Hallahan, N. A. Cogliano-Shutta, J. A. Metcalf, M. McLaughlin, R. Kwan, J. M. Mican, R. T. Davey, M. Connors. Defective human immunodeficiency virus-specific CD8+ T-cell polyfunctionality, proliferation, and cytotoxicity are not restored by antiretroviral therapy. J. Virol. 83, 11876–11889 (2009).

23. Z. M. Ndhlovu, L. B. Chibnik, J. Proudfoot, S. Vine, A. McMullen, K. Cesa, F. Porichis, B. R. Jones, D. Alvino, M. G. Hart, E. Stampouloglou, A. Piechocka-Trocha, C. Kadie, F. Pereyra, D. Heckerman, P. L. Jager, B. D. Walker, D. E. Kaufmann. High-dimensional immunomonitoring models of HIV-1-specific CD8 T-cell responses accurately identify subjects achieving spontaneous viral control. Blood. 121, 801–811 (2013).

24. Z. M. Ndhlovu, J. Proudfoot, K. Cesa, D. Alvino, A. McMullen, S. Vine, E. Stampouloglou, Piechocka-Trocha, B. D. Walker, F. Pereyra. Elite controllers with low to absent effector CD8+ T cell responses maintain highly functional, broadly directed central memory responses. J. Virol. 86, 6959–6969 (2012).

25. T. Yamamoto, D. A. Price, J. P. Casazza, G. Ferrari, M. Nason, P. K. Chattopadhyay, M. Roederer, E. Gostick, P. D. Katsikis, D. C. Douek, R. Haubrich, C. Petrovas, R. A. Koup. Surface expression patterns of negative regulatory molecules identify determinants of virus-specific CD8+ T-cell exhaustion in HIV infection. Blood. 117, 4805–4815 (2011).

26. S. Viganò, R. Banga, F. Bellanger, C. Pellaton, A. Farina, D. Comte, A. Harari, M. Perreau. CD160-associated CD8 T-cell functional impairment is independent of PD-1 expression. PLoS Pathog. 10, e1004380 (2014).

27. C. Pombo, J. E. Wherry, E. Gostick, D. A. Price, M. R. Betts. Elevated Expression of CD160 and 2B4 Defines a Cytolytic HIV-Specific CD8+ T-Cell Population in Elite Controllers. J. Infect. Dis. 212, 1376–1386 (2015).

28. G. D. Gaiha, K. J. McKim, M. Woods, T. Pertel, J. Rohrbach, N. Barteneva, C. R. Chin, D. Liu, D. Z. Soghoian, K. Cesa, S. Wilton, M. T. Waring, A. Chicoine, T. Doering, J. E. Wherry, D. E. Kaufmann, M. Lichterfeld, A. L. Brass, B. D. Walker. Dysfunctional HIV-specific CD8+ T cell proliferation is associated with increased caspase-8 activity and mediated by necroptosis. Immunity. 41, 1001–1012 (2014).

29. M. Quigley, F. Pereyra, B. Nilsson, F. Porichis, C. Fonseca, Q. Eichbaum, B. Julg, J. L. Jesneck, K. Brosnahan, S. Imam, K. Russell, I. Toth, A. Piechocka-Trocha, D. Dolfi, J. Angelosanto, A. Crawford, H. Shin, D. S. Kwon, J. Zupkosky, L. Francisco, G. J. Freeman, J. E. Wherry, D. E. Kaufmann, B. D. Walker, B. Ebert, N. W. Haining. Transcriptional analysis of HIV-specific CD8+ T cells shows that PD-1 inhibits T cell function by upregulating BATF. Nat. Med. 16, 1147–1151 (2010).

30. S. Verbeek, D. Izon, F. Hofhuis, E. Robanus-Maandag, H. te Riele, M. van de Wetering, M. Oosterwegel, A. Wilson, H. MacDonald, H. Clevers. An HMG-box-containing T-cell factor required for thymocyte differentiation. Nature. 374, 70–74 (1995).

31. G. Jeannet, C. Boudousquié, N. Gardiol, J. Kang, J. Huelsken, W. Held. Essential role of the Wnt pathway effector Tcf-1 for the establishment of functional CD8 T cell memory. P. Natl. A.Sci. 107, 9777–9782 (2010).

32. X. Zhou, S. Yu, D.-M. Zhao, J. T. Harty, V. P. Badovinac, H.-H. Xue. Differentiation and persistence of memory CD8(+) T cells depend on T cell factor 1. Immunity. 33, 229–240 (2010).

33. D.-M. Zhao, S. Yu, X. Zhou, J. S. Haring, W. Held, V. P. Badovinac, J. T. Harty, H.-H. Xue. Constitutive activation of Wnt signaling favors generation of memory CD8 T cells. J. Immunol. 184, 1191–1199 (2010).

34. R. Kratchmarov, A. M. Magun, S. L. Reiner. TCF1 expression marks self-renewing human CD8+ T cells. Blood Adv. 2, 1685–1690 (2018).

35. X. Zhou, H.-H. Xue, Cutting edge: generation of memory precursors and functional memory CD8+ T cells depends on T cell factor-1 and lymphoid enhancer-binding factor-1. J. Immunol. 189, 2722–2726 (2012).

36. T. Wu, Y. Ji, A. E. Moseman, H. C. Xu, M. Manglani, M. Kirby, S. M. Anderson, R. Handon, E. Kenyon, A. Elkahloun, W. Wu, P. A. Lang, L. Gattinoni, D. B. McGavern, P. L. Schwartzberg. The TCF1-Bcl6 axis counteracts type I interferon to repress exhaustion and maintain T cell stemness. Sci. Immunol. 1, eaai8593 (2016).

37. D. T. Utzschneider, M. Charmoy, V. Chennupati, L. Pousse, D. Ferreira, S. Calderon-Copete, M. Danilo, F. Alfei, M. Hofmann, D. Wieland, S. Pradervand, R. Thimme, D. Zehn, W. Held. T Cell Factor 1-Expressing Memory-like CD8(+) T Cells Sustain the Immune Response to Chronic Viral Infections. Immunity. 45, 415–427 (2016).

38. S. Im, M. Hashimoto, M. Y. Gerner, J. Lee, H. T. Kissick, M. C. Burger, Q. Shan, S. J. Hale, J. Lee, T. H. Nasti, A. H. Sharpe, G. J. Freeman, R. N. Germain, H. I. Nakaya, H.-H. Xue, R. Ahmed. Defining CD8(+) T cells that provide the proliferative burst after PD-1 therapy. Nature. 537, 417–421 (2016).

39. A. Schuch, E. Alizei, K. Heim, D. Wieland, M. Kiraithe, J. Kemming, S. Llewellyn-Lacey, Ö. Sogukpinar, Y. Ni, S. Urban, P. Zimmermann, M. Nassal, F. Emmerich, D. A. Price, B. Bengsch, H. Luxenburger, C. Neumann-Haefelin, M. Hofmann, R. Thimme. Phenotypic and functional differences of HBV core-specific versus HBV polymerase-specific CD8+ T cells in chronically HBV-infected patients with low viral load. Gut. 68, 905–915 (2019).

40. H. Kefalakes, C. Koh, J. Sidney, G. Amanakis, A. Sette, T. Heller, B. Rehermann. Hepatitis D Virus-Specific CD8+ T Cells Have a Memory-Like Phenotype Associated With Viral Immune Escape in Patients With Chronic Hepatitis D Virus Infection. Gastroenterology. 156, 1805–1819.e9 (2019).

41. D. Wieland, J. Kemming, A. Schuch, F. Emmerich, P. Knolle, C. Neumann-Haefelin, W. Held, D. Zehn, M. Hofmann, R. Thimme. TCF1(+) hepatitis C virus-specific CD8(+) T cells are maintained after cessation of chronic antigen stimulation. Nat. Commun. 8, 15050 (2017).

42. M. Sade-Feldman, K. Yizhak, S. L. Bjorgaard, J. P. Ray, C. G. de Boer, R. W. Jenkins, D. J. Lieb, J. H. Chen, D. T. Frederick, M. Barzily-Rokni, S. S. Freeman, A. Reuben, P. J. Hoover, A.-C. Villani, E. Ivanova, A. Portell, P. H. Lizotte, A. R. Aref, J.-P. Eliane, M. R. Hammond, H. Vitzthum, S. M. Blackmon, B. Li, V. Gopalakrishnan, S. M. Reddy, Z. A. Cooper, C. P. Paweletz, D. A. Barbie, A. Stemmer-Rachamimov, K. T. Flaherty, J. A. Wargo, G. M. Boland, R. J. Sullivan, G. Getz, N. Hacohen. Defining T Cell States Associated with Response to Checkpoint Immunotherapy in Melanoma. Cell. 175, 998–1013.e20 (2018).

43. B. C. Miller, D. R. Sen, R. Abosy, K. Bi, Y. V. Virkud, M. W. LaFleur, K. B. Yates, A. Lako, K. Felt, G. S. Naik, M. Manos, E. Gjini, J. R. Kuchroo, J. J. Ishizuka, J. L. Collier, G. K. Griffin, S. Maleri, D. E. Comstock, S. A. Weiss, F. D. Brown, A. Panda, M. D. Zimmer, R. T. Manguso, S. F. Hodi, S. J. Rodig, A. H. Sharpe, N. W. Haining. Subsets of exhausted CD8+ T cells differentially mediate tumor control and respond to checkpoint blockade. Nat. Immunol. 20, 326–336 (2019).

44. Z. Chen, Z. Ji, S. Ngiow, S. Manne, Z. Cai, A. C. Huang, J. Johnson, R. P. Staupe, B. Bengsch, C. Xu, S. Yu, M. Kurachi, R. S. Herati, L. A. Vella, A. E. Baxter, J. E. Wu, O. Khan, J.-C. Beltra, J. R. Giles, E. Stelekati, L. M. McLane, C. Lau, X. Yang, S. L. Berger, G. Vahedi, H. Ji, J. E. Wherry. TCF-1-Centered Transcriptional Network Drives an Effector versus Exhausted CD8 T Cell-Fate Decision. Immunity. 51, 840–855 (2019).

45. Y. Leong, Y. Chen, H. Ong, D. Wu, K. Man, C. Deleage, M. Minnich, B. J. Meckiff, Y. Wei, Z. Hou, D. Zotos, K. A. Fenix, A. Atnerkar, S. Preston, J. G. Chipman, G. J. Beilman, C. C. Allison, L. Sun, P. Wang, J. Xu, J. G. Toe, H. K. Lu, Y. Tao, U. Palendira, A. L. Dent, A. L. Landay, M. Pellegrini, I. Comerford, S. R. McColl, T. W. Schacker, H. M. Long, J. D. Estes, M. Busslinger, G. T. Belz, S. R. Lewin, A. Kallies, D. Yu. CXCR5(+) follicular cytotoxic T cells control viral infection in B cell follicles. Nat. Immunol. 17, 1187–1196 (2016).

46. S. Viganò, J. Negron, Z. Ouyang, E. S. Rosenberg, B. D. Walker, M. Lichterfeld, X. G. Yu, Prolonged Antiretroviral Therapy Preserves HIV-1-Specific CD8 T Cells with Stem Cell-Like Properties. J. Virol. 89, 7829–7840 (2015).

47. D. Mendoza, S. A. Migueles, J. E. Rood, B. Peterson, S. Johnson, N. Doria-Rose, D. Schneider, E. Rakasz, M. T. Trivett, C. M. Trubey, V. Coalter, C. W. Hallahan, D. Watkins, G. Franchini, J. D. Lifson, M. Connors. Cytotoxic Capacity of SIV-Specific CD8+ T Cells against Primary Autologous Targets Correlates with Immune Control in SIV-Infected Rhesus Macaques. Plos Pathog. 9, e1003195 (2013).

48. N. Takemoto, A. M. Intlekofer, J. T. Northrup, J. E. Wherry, S. L. Reiner. Cutting Edge: IL-12 Inversely Regulates T-bet and Eomesodermin Expression during Pathogen-Induced CD8 + T Cell Differentiation. J. Immunol. 177, 7515–7519 (2006).

49. M. Paley, D. Kroy, P. Odorizzi, J. Johnnidis, D. Dolfi, B. Barnett, E. Bikoff, E. Robertson, G. Lauer, S. Reiner, E. Wherry. Progenitor and terminal subsets of CD8+ T cells cooperate to contain chronic viral infection. Science. 338, 1220–1225 (2012).

50. R. M. Okamura, M. Sigvardsson, J. Galceran, S. Verbeek, H. Clevers, R. Grosschedl. Redundant Regulation of T Cell Differentiation and TCRα Gene Expression by the Transcription Factors LEF-1 and TCF-1. Immunity. 8, 11–20 (1998).

51. T. Willinger, T. Freeman, M. Herbert, H. Hasegawa, A. J. McMichael, M. F. Callan, Human naive CD8 T cells down-regulate expression of the WNT pathway transcription factors lymphoid enhancer binding factor 1 and transcription factor 7 (T cell factor-1) following antigen encounter in vitro and in vivo. J. Immunol. 176, 1439–1446 (2006).

52. J. Hendriks, Y. Xiao, J. Borst. CD27 promotes survival of activated T cells and complements CD28 in generation and establishment of the effector T cell pool. J. Exp. Med. 198, 1369–80 (2003).

53. A. F. Ochsenbein, S. R. Riddell, M. Brown, L. Corey, G. M. Baerlocher, P. M. Lansdorp, P. D. Greenberg. CD27 Expression Promotes Long-Term Survival of Functional Effector–Memory CD8+Cytotoxic T Lymphocytes in HIV-infected Patients. J. Exp. Med. 200, 1407–1417 (2004).

54. G. M. Chew, T. Fujita, G. M. Webb, B. J. Burwitz, H. L. Wu, J. S. Reed, K. B. Hammond, K. L. Clayton, N. Ishii, M. Abdel-Mohsen, T. Liegler, B. I. Mitchell, F. M. Hecht, M. Ostrowski, C. ikuma, S. G. Hansen, M. Maurer, A. J. Korman, S. G. Deeks, J. B. Sacha, L. C. Ndhlovu. TIGIT Marks Exhausted T Cells, Correlates with Disease Progression, and Serves as a Target for Immune Restoration in HIV and SIV Infection. PLoS Pathog. 12, e1005349 (2016).

55. Y. Peretz, Z. He, Y. Shi, B. Yassine-Diab, J.-P. Goulet, R. Bordi, A. Filali-Mouhim, J.-B. Loubert, M. El-Far, F. P. Dupuy, M.-R. Boulassel, C. Tremblay, J.-P. Routy, N. Bernard, R. Balderas, E. K. Haddad, R.-P. Sékaly. CD160 and PD-1 co-expression on HIV-specific CD8 T cells defines a subset with advanced dysfunction. PLoS Pathog. 8, e1002840 (2012).

56. V. K. Kuchroo, A. C. Anderson, C. Petrovas. Coinhibitory receptors and CD8 T cell exhaustion in chronic infections. Curr. Opin. HIV AIDS. 9, 439–445 (2014).

57. C. M. Ramirez, E. Sinclair, L. Epling, S. A. Lee, V. Jain, P. Y. Hsue, H. Hatano, D. Conn, F. M. Hecht, J. N. Martin, J. M. McCune, S. G. Deeks, P. W. Hunt. Immunologic profiles distinguish aviremic HIV-infected adults. AIDS. 30, 1553–62 (2016).

58. T. L. Roth, J. P. Li, J. F. Nies, R. Yu, M. L. Nguyen, Y. Lee, R. Apathy, A. Truong, J. Hiatt, D. Wu, D. N. Nguyen, D. Goodman, J. A. Bluestone, K. Roybal, E. Shifrut, A. Marson. Rapid discovery of synthetic DNA sequences to rewrite endogenous T cell circuits. bioRxiv. 12, doi: https://doi.org/10.1101/604561 (2019).

59. T. L. Roth, C. Puig-Saus, R. Yu, E. Shifrut, J. Carnevale, J. P. Li, J. Hiatt, J. Saco, P. Krystofinski, H. Li, V. Tobin, D. N. Nguyen, M. R. Lee, A. L. Putnam, A. L. Ferris, J. W. Chen, J.-N. Schickel, L. Pellerin, D. Carmody, G. Alkorta-Aranburu, D. Gaudio, H. Matsumoto, M. Morell, Y. Mao, M. Cho, R. M. Quadros, C. B. Gurumurthy, B. Smith, M. Haugwitz, S. H. Hughes, J. S. Weissman, K. Schumann, J. H. Esensten, A. P. May, A. Ashworth, G. M. Kupfer, S. W. Greeley, R. Bacchetta, E. Meffre, M. Roncarolo, N. Romberg, K. C. Herold, A. Ribas, M. D. Leonetti, A. Marson. Reprogramming human T cell function and specificity with non-viral genome targeting. Nature. 559, 405–409 (2018).

60. A. Sharma, Q. Chen, T. Nguyen, Q. Yu, J. Sen. T cell factor-1 and β-catenin control the development of memory-like CD8 thymocytes. J. Immunol. 188, 3859–3868 (2012).

61. D. Raghu, H.-H. Xue, L. A. Mielke. Control of Lymphocyte Fate, Infection, and Tumor Immunity by TCF-1. Trends Immunol. 40, 1149–1162 (2019).

62. M. M. Tiemessen, M. R. Baert, L. Kok, M. C. van Eggermond, P. J. van den Elsen, R. Arens, F. J. Staal. T Cell factor 1 represses CD8+ effector T cell formation and function. J. Immunol. 193, 5480–5487 (2014).

63. I. Siddiqui, K. Schaeuble, V. Chennupati, S. A. Marraco, S. Calderon-Copete, D. Ferreira, S. J. Carmona, L. Scarpellino, D. Gfeller, S. Pradervand, S. A. Luther, D. E. Speiser, W. Held, Intratumoral Tcf1+PD-1+CD8+ T Cells with Stem-like Properties Promote Tumor Control in Response to Vaccination and Checkpoint Blockade Immunotherapy. Immunity. 50, 195–211. (2019).

64. W. Held, I. Siddiqui, K. Schaeuble, D. E. Speiser. Intratumoral CD8+ T cells with stem cell-like properties: Implications for cancer immunotherapy. Sci. Transl. Med. 11, eaay6863 (2019).

65. B. Liu, W. Zhang, H. Zhang. Development of CAR-T cells for long-term eradication and surveillance of HIV-1 reservoir. Curr. Opin. Virol. 38, 21–30 (2019).

66. A. D. McLellan, S. M. Rad. Chimeric antigen receptor T cell persistence and memory cell formation. Immunol. Cell Bio. 97, 664–674 (2019).

67. L. Gattinoni, X.-S. Zhong, D. C. Palmer, Y. Ji, C. S. Hinrichs, Z. Yu, C. Wrzesinski, A. Boni, L. Cassard, L. M. Garvin, C. M. Paulos, P. Muranski, N. P. Restifo. Wnt signaling arrests effector T cell differentiation and generates CD8+ memory stem cells. Nat. Med. 15, 808–813 (2009).

68. E. Connick, J. M. Folkvord, K. T. Lind, E. G. Rakasz, B. Miles, N. A. Wilson, M. L. Santiago, K. Schmitt, E. B. Stephens, H. O. Kim, R. Wagstaff, S. Li, H. M. Abdelaal, N. Kemp, D. I. Watkins, S. MaWhinney, P. J. Skinner. Compartmentalization of simian immunodeficiency virus replication within secondary lymphoid tissues of rhesus macaques is linked to disease stage and inversely related to localization of virus-specific CTL. J. Immunol. 193, 5613–5625 (2014).

69. M. P. Bronnimann, P. J. Skinner, E. Connick. The B-Cell Follicle in HIV Infection: Barrier to a Cure. Front. Immunol. 9, 20 (2018).

70. S. Pallikkuth, M. Sharkey, D. Z. Babic, S. Gupta, G. W. Stone, M. A. Fischl, M. Stevenson, S. Pahwa. Peripheral T Follicular Helper Cells Are the Major HIV Reservoir within Central Memory CD4 T Cells in Peripheral Blood from Chronically HIV-Infected Individuals on Combination Antiretroviral Therapy. J. Virol. 90, 2718–28 (2015).

71. M. Buggert, S. Nguyen, G. de Oca, B. Bengsch, S. Darko, A. Ransier, E. R. Roberts, D. Alcazar, I. Brody, L. A. Vella, L. Beura, S. Wijeyesinghe, R. S. Herati, P. l Estrada, Y. Ablanedo-Terrazas, L. Kuri-Cervantes, A. Japp, S. Manne, S. Vartanian, A. Huffman, J. K. Sandberg, E. Gostick, G. Nadolski, G. Silvestri, D. H. Canaday, D. A. Price, C. Petrovas, L. F. Su, G. Vahedi, Y. Dori, I. Frank, M. G. Itkin, J. E. Wherry, S. G. Deeks, A. Naji, G. Reyes-Terán, D. Masopust, D. C. Douek, M. R. Betts. Identification and characterization of HIV-specific resident memory CD8+ T cells in human lymphoid tissue. Sci. Immunol. 3, eaar4526 (2018).

72. S. Nguyen, C. Deleage, S. Darko, A. Ransier, D. P. Truong, D. Agarwal, A. Japp, V. H. Wu, Kuri-Cervantes, M. Abdel-Mohsen, P. l Estrada, Y. Ablanedo-Terrazas, E. Gostick, J. A. Hoxie, N. R. Zhang, A. Naji, G. Reyes-Terán, J. D. Estes, D. A. Price, D. C. Douek, S. G. Deeks, M. Buggert, M. R. Betts. Elite control of HIV is associated with distinct functional and transcriptional signatures in lymphoid tissue CD8+ T cells. Sci. Transl. Med. 11, eaax4077 (2019).

73. F. Perdomo-Celis, N. Taborda, M. Rugeles. Follicular CD8+ T Cells: Origin, Function and Importance during HIV Infection. Front. Immunol. 8, 1241 (2017).

74. S. Li, J. M. Folkvord, E. G. Rakasz, H. M. Abdelaal, R. K. Wagstaff, K. J. Kovacs, H. O. Kim, R. Sawahata, S. MaWhinney, D. Masopust, E. Connick, P. J. Skinner. Simian Immunodeficiency Virus-Producing Cells in Follicles Are Partially Suppressed by CD8+ Cells In Vivo. J. Virol. 90, 11168–11180 (2016).

75. F. M. Behr, N. A. Kragten, T. H. Wesselink, B. Nota, R. A. van Lier, D. Amsen, R. Stark, P. Hombrink, K. P. van Gisbergen. Blimp-1 Rather Than Hobit Drives the Formation of Tissue-Resident Memory CD8+ T Cells in the Lungs. Front. Immunol. 10, 400 (2019).

76. M. A. Reuter, P. l Estrada, M. Buggert, C. Petrovas, S. Ferrando-Martinez, S. Nguyen, A. Japp, Y. Ablanedo-Terrazas, A. Rivero-Arrieta, L. Kuri-Cervantes, H. M. Gunzelman, E. Gostick, D. A. Price, R. A. Koup, A. Naji, D. H. Canaday, G. Reyes-Terán, M. R. Betts. HIV-Specific CD8 + T Cells Exhibit Reduced and Differentially Regulated Cytolytic Activity in Lymphoid Tissue. Cell Rep. 21, 3458–3470 (2017).

77. B. Kiniry, A. Ganesh, J. Critchfield, P. Hunt, F. Hecht, msouk, S. Deeks, B. Shacklett. Predominance of weakly cytotoxic, T-bet(Low)Eomes(Neg) CD8(+) T-cells in human gastrointestinal mucosa: implications for HIV infection. Mucosal Immunol. 10, 1008–1020 (2016).

78. W. J. Critchfield, D. H. Young, T. L. Hayes, J. V. Braun, J. C. Garcia, R. B. Pollard, B. L. Shacklett. Magnitude and complexity of rectal mucosa HIV-1-specific CD8+ T-cell responses during chronic infection reflect clinical status. PLoS ONE. 3, e3577 (2008).

79. I. J. Fuss, M. E. Kanof, P. D. Smith, H. Zola. Isolation of whole mononuclear cells from peripheral blood and cord blood. Curr Protoc Immunol. Chapter 7, Unit7.1 (2001).

80. R. E. Owen, E. Sinclair, B. Emu, J. W. Heitman, D. F. Hirschkorn, L. C. Epling, Q. Tan, B. Custer, J. M. Harris, M. A. Jacobson, J. M. McCune, J. N. Martin, F. M. Hecht, S. G. Deeks, P. J. Norris. Loss of T cell responses following long-term cryopreservation. J. Immunol. Methods. 326, 93–115 (2007).

81. C. E. Starke, C. L. Vinton, K. Ladell, J. E. McLaren, A. M. Ortiz, J. C. Mudd, J. K. Flynn, S. H. Lai, F. Wu, V. M. Hirsch, S. Darko, D. C. Douek, D. A. Price, J. M. Brenchley. SIV-specific CD8+ T cells are clonotypically distinct across lymphoid and mucosal tissues. J. Clin. Invest. doi:10.1172/jci129161 (2019).

82. C. T. Deguit, M. Hough, R. Hoh, M. Krone, C. D. Pilcher, J. N. Martin, S. G. Deeks, J. M. McCune, P. W. Hunt, R. L. Rutishauser. Some Aspects of CD8+ T Cell Exhaustion are Associated with Altered T Cell Mitochondrial Features and ROS content in HIV Infection. J. Acquir. Defic. Syndr. 82, 211–219 (2019).

83. NIH Tetramer Core Facility. Class I MHC Tetramer Preparation: Overview. Production Protocols Web Site. Emory University. https://tetramer.yerkes.emory.edu/support/protocols (2005).

84. J. F. Hultquist, J. Hiatt, K. Schumann, M. J. McGregor, T. L. Roth, P. Haas, J. A. Doudna, A. Marson, N. J. Krogan. CRISPR-Cas9 genome engineering of primary CD4+ T cells for the interrogation of HIV-host factor interactions. Nat. Protoc. 14, 1–27 (2019).

85. A. Pasetto, A. Gros, P. F. Robbins, D. C. Deniger, T. D. Prickett, R. Matus-Nicodemos, D. C. Douek, B. Howie, H. Robins, M. R. Parkhurst, J. Gartner, K. Trebska-McGowan, J. S. Crystal, S. A. Rosenberg. Tumor- and Neoantigen-Reactive T-cell Receptors Can Be Identified Based on Their Frequency in Fresh Tumor. Cancer Immunol. Res. 4, 734–43 (2016).

86. D. N. Nguyen, T. L. Roth, J. P. Li, P. Chen, R. Apathy, M. R. Mamedov, L. T. Vo, V. R. Tobin, D. Goodman, E. Shifrut, J. A. Bluestone, J. M. Puck, F. C. Szoka, A. Marson. Polymer-stabilized Cas9 nanoparticles and modified repair templates increase genome editing efficiency. Nat. Biotechnol. 1–6 (2019).

